# Preserved implicit metacognitive sensitivity distinguishes psychosis risk from first-episode psychosis: A cross-sectional virtual reality-based study

**DOI:** 10.64898/2026.08.09.743764

**Authors:** Yonatan Stern, Donia Sussan, Barnaby Nelson, Uri Hertz, Morris Goldsmith, Eyal Bergmann, Lauren Nashashibi, Roy Salomon, Danny Koren

## Abstract

Relatively preserved insight distinguishes individuals at risk for psychosis from those with full-blown psychosis. Metacognitive processes thought to support insight and uncertainty monitoring may therefore serve as early markers of illness progression. Yet findings have been inconsistent, perhaps partly due to reliance on explicit confidence ratings that introduce reflection and response biases. To address these limitations, we used a novel implicit confidence measure derived from post-decision gaze in a virtual-reality probabilistic learning task. Gaze-based confidence quantifies the alignment between spatial predictions and gaze direction. We assessed first-order learning and gaze-based metacognition in four groups: clinical high-risk for psychosis (CHR-P), first episode psychosis (FEP), help-seeking controls (HSC), and healthy controls (HC). We tested whether implicit metacognition differentiates psychosis risk from psychosis.

Learning accuracy was reduced in both CHR-P and FEP compared to control groups. At the metacognitive level, CHR-P confidence levels were approximately commensurate with their reduced first-order performance, indicating preserved confidence calibration, along with preserved metacognitive sensitivity—the ability to distinguish correct from incorrect decisions. In contrast, FEP showed impaired confidence calibration and reduced metacognitive sensitivity. Metacognitive calibration and sensitivity distinguished CHR-P from FEP and provided predictive value in distinguishing CHR-P from FEP, whereas learning accuracy did not.

These findings reveal a dissociation between first-order cognitive processes and distinct aspects of implicit metacognition, including confidence calibration and metacognitive sensitivity, across the psychosis continuum. This dissociation may refine early clinical characterization and improve identification of preserved insight-related mechanisms in psychosis risk.

## 1. Introduction

### 1.1 Metacognition in clinical versus sub-clinical psychosis

Impaired insight into the anomalous nature of one’s experiences is a defining feature of psychosis (Amador et al., 1994). In contrast, individuals at clinical high risk for psychosis (CHR-P) retain the capacity to question the veridicality of their experiences, making preserved insight a key distinguishing diagnostic criterion (McGlashan et al., 2001; Miller et al., 2003). Accordingly, growing interest has focused on the cognitive processes that underlie insight, particularly the evaluation of uncertainty in one’s own perceptions and decisions. One such process is metacognition—the ability to monitor and evaluate one’s cognitive processes and the reliability of one’s judgments (Flavell, 1979; Nelson, 1990).

Two complementary aspects of metacognitive monitoring are particularly relevant: calibration and sensitivity. Metacognitive calibration reflects the correspondence between overall subjective confidence and overall objective performance, whereas metacognitive sensitivity reflects the extent to which confidence discriminates between correct and incorrect decisions. (Maniscalco and Lau, 2012; Rahnev, D., 2023). Reduced metacognitive calibration and sensitivity have been documented in psychosis (Koren et al., 2004, 2005; Krugwasser et al., 2021; for review see Hohendorf and Bauer, 2023 but see also Rouy et al., 2021) and in individuals with genetic liability for psychosis (Salomon et al., 2021). Whereas, altered calibration has been associated with schizotypal traits in the general population (Lehmann and Ettinger, 2023; Rouault et al., 2018).

Despite the close conceptual link between metacognition and insight, findings regarding metacognitive sensitivity during the psychosis-risk phase remain inconsistent. While some studies have reported metacognitive performance in psychosis-risk populations that is comparable to healthy controls and superior to that observed in full-blown psychosis (Eisenacher et al., 2015; Gawęda et al., 2018), others have not replicated these findings (Köther et al., 2018).

### 1.2 Implicit measures of confidence as a possible solution

The inconsistent findings regarding metacognitive sensitivity in psychosis-risk populations may partly reflect limitations of current methods of metacognitive assessment. Most previous studies have relied on explicit confidence ratings obtained after each decision. Such reports may alter the very processes they are intended to measure by inducing reflection (Double and Birney, 2019) and introducing response biases that are associated with psychiatric symptoms (Sarna et al., 2026). Consequently, explicit confidence ratings may obscure spontaneous uncertainty monitoring and thereby reduce sensitivity to early alterations that distinguish psychosis risk from full-blown psychosis.

Internal confidence is also reflected in spontaneous behavioral and physiological signals, including response times (Kiani et al., 2014) and sensorimotor adaptation (Faivre et al., 2020). These manifestations provide a more direct window onto uncertainty monitoring because they neither require explicit introspection nor interrupt task performance. The present study therefore examined an implicit behavioral manifestation of confidence derived from post-decision gaze.

### 1.3 Gaze-based measurement of metacognition: Empirical validation

Among implicit behavioral measures, gaze is particularly relevant in psychosis research, as eye-movement abnormalities represent one of the most robust behavioral markers across the psychosis spectrum (Benson et al., 2012; Thakkar et al., 2017), reflecting a decoupling of attention from learned expectations and prior knowledge (Adámek et al., 2024; Miura et al., 2025). Building on this work, our group recently developed and validated a novel implicit measure of confidence based on post-decision gaze behavior (Stern et al., 2026).

Participants completed a perceptual decision-making task in which they selected one of two briefly presented stimuli. During a subsequent post-decision interval, gaze behavior was continuously recorded while no explicit confidence judgments were required. Confidence was inferred from the relative time participants spontaneously viewed the chosen direction versus unchosen option, with greater gaze toward the selected option indicating higher confidence in the decision.

Converging evidence supported the validity of this measure as an implicit behavioral manifestation of confidence (Stern et al., 2026). It predicted explicit confidence ratings, tracked trial-by-trial variation in computationally derived decision certainty, and exhibited statistical hallmarks of confidence (Sanders et al., 2016). Together, these findings suggest that post-decision gaze provides a valid implicit behavioral manifestation of internal confidence, making it well suited for investigating uncertainty monitoring across the psychosis continuum.

### 1.4 The present study: Goals and hypotheses

The present study used a virtual reality-based probabilistic learning task with eye tracking to examine implicit metacognitive functioning across the psychosis continuum. By jointly assessing first-order learning and gaze-based confidence, we tested whether confidence calibration and metacognitive sensitivity distinguish CHR-P from FEP beyond differences in first-order performance.

The study hypotheses were preregistered (osf.io/bc9np/overview) and refined in light of theoretical and empirical developments that emerged after preregistration (see Supplementary Materials [SM]). The following hypotheses were tested:

#### H1

Based on evidence of disrupted predictive processing (Katthagen et al., 2022) and overestimation of the environment’s volatility in early psychosis (Cole et al., 2020), we hypothesized that first-order learning accuracy would be reduced in both CHR-P and FEP relative to controls. We further hypothesized that they would exhibit an increased tendency to switch predictions following correct predictions (a “Win-Switch” strategy).

#### H2

Consistent with the relatively preserved insight that characterizes CHR-P (McGlashan et al., 2001; Miller et al., 2003), we hypothesized that confidence calibration would differ across the psychosis continuum. Specifically, we hypothesized that CHR-P participants would exhibit reduced gaze-based confidence relative to HC that is commensurate with their reduced first-order performance, whereas FEP participants would not show a corresponding reduction in confidence despite impaired performance. We further hypothesized that CHR-P participants would continue to exhibit lower confidence than HC after adjusting for first-order performance.

#### H3a

Consistent with CHR-P’s relatively preserved insight, we hypothesized that gaze-based metacognitive sensitivity would be preserved in the CHR-P group but impaired in the FEP group.

#### H3b

We further hypothesized that implicit metacognitive sensitivity would constitute the strongest single predictor distinguishing CHR-P from FEP group membership and provide significant incremental predictive value beyond first-order performance.

Together, these hypotheses test whether first-order learning and implicit metacognitive monitoring become dissociated across the psychosis continuum, thereby distinguishing psychosis risk from first-episode psychosis.

## 2. Methods

### 2.1 Participants

Participants were 18–35 years old, right-handed, with normal or corrected-to-normal vision and no history of serious head injury. All participants provided written informed consent and were compensated 50 NIS per hour. The study received approval from the Internal Review Board of the School of Psychological Sciences, University of Haifa (Approval # 072/23), and the Rambam Medical Center Helsinki Committee (Study 0718-19 RMB).

### 2.3 Clinical Measures

#### 2.3.1 Clinical assessments

Clinical status was determined using semi-structured diagnostic interviews administered by trained clinical psychologists under regular supervision (D.K.).

Participants completed: (1) the Structured Interview for Psychosis-Risk Syndromes (SIPS), to assess CHR-P criteria and severity of attenuated psychotic symptoms (McGlashan et al., 2001); (2) the Brief Psychiatric Rating Scale (BPRS), to assess general psychiatric symptoms (Overall and Gorham, 1988); and (3) the Mini International Neuropsychiatric Interview (MINI), to establish DSM-based psychiatric diagnoses (Sheehan et al., 1998), corroborated by medical records when available. Participants also completed semi-structured interviews (i.e., the Examination of Anomalous Self-Experiences [EASE]; (Parnas et al., 2005) to evaluate self-disturbances) and self-report questionnaires (see SM for details) for exploratory analyses outside the scope of the present report.

#### 2.3.2 Clinical group assignment

Following the preregistered criteria, participants were assigned to one of four mutually exclusive groups (see Table 1 for characteristics): (1) Healthy Control (HC) group (*N* = 45; 20 male) comprising individuals with no psychiatric diagnosis, no use of psychotropic medication, and did not seek or receive mental-health services in the past year (2) Help-Seeking Controls (HSC) group (*N* = 47; 13 male) comprising individuals that sought mental-health services and met criteria for a non-psychotic disorder (see SM for clinical diagnosis) but did not meet CHR-P criteria, (3) Clinical High-Risk for Psychosis (CHR-P) group (*N* = 31; 21 male) comprising individuals who met SIPS criteria for an at-risk syndrome (27 with attenuated psychosis syndrome, 2 with brief limited intermittent psychotic symptoms, and 2 with genetic risk and deterioration), (4) First-Episode Psychosis (FEP) group (*N* = 21; 13 male) comprising individuals who experienced a full psychotic episode within the past three years (8 with schizoaffective disorder, 6 with schizophrenia, 6 with psychosis not otherwise specified, and one with bipolar).

**Table 1.** Socio-demographic and clinical characteristics of participant groups.

|  | 1. Healthy Controls (HC)<br>N=45 | 2. Help-Seeking Controls (HSC)<br>N=47 | 3. Clinical High Risk for Psychosis (CHR-P)<br>N=31 | 4. First Episode of Psychosis (FEP)<br>N=21 | Significance test | Post-hoc Tukey test |
| --- | --- | --- | --- | --- | --- | --- |
| Age (years)<br>(Mean [95% CI]) | 23.8<br>[22.9, 24.7] | 25.7<br>[25.1, 26.3] | 23.6<br>[22.2, 25.0] | 26.1<br>[24.2, 28.0] | $F_{(4.2, 140)} = 4.2, p = .01$ | 1~2~3~4 |
| Gender (N<sub>male</sub>) | 20 | 14 | 21 | 14 | $\chi^2_{(3)} = 14.3, p = .003$ | 1~2<3~4 |
| Mean years of education | 12.8<br>[12.4, 13.2] | 13<br>[12.4, 13.6] | 12.5<br>[12.0, 13.0] | 13.1<br>[12.6, 13.6] | $F_{(0.76, 117)} < 1, p = .52$ | ---- |
| GAF<br>(Mean [95% CI]) | 86.2<br>[83.5, 88.9] | 64.2<br>[60.9, 67.5] | 51.3<br>[46.6, 56.0] | 52.5 [47.5, 57.5] | $F_{(3, 131)} = 81.9, p < .001$ | 1<2<3~4 |
| Previously used VR (%) | 30 | 32.5 | 29.6 | 38.9 | $\chi^2_{(3)} = 0.5, p = .9$ | ---- |
| Took psychiatric medication in past year (% yes) | 0 | 61 | 61.3 | 81 | $\chi^2_{(3)} = 59, p < .001$ | 1<2~3~4 |
| In therapy in past year (% yes) | 4.4 | 57.4 | 71 | 61.9 | $\chi^2_{(3)} = 43.8, p < .001$ | 1<(2~3~4) |
| Has been in psychiatric hospitalization (% yes) | 0 | 25.5 | 32.3 | 85.7 | $\chi^2_{(3)} = 49.4, p < .001$ | 1<(2~3)<4 |
| Diagnosed with ADHD (% yes) | 17.8 | 44.7 | 28.6 | 60.0 | $\chi^2_{(3)} = 13.8, p < .001$ | 1<4~2~3 |
| Family history of mental health diagnosis (% yes) | 11.1 | 45.7 | 25 | 33.3 | $\chi^2_{(3)} = 13.7, p < .001$ | (1<2)~3~4 |
| Sum of SIPS positive subscales (Mean [95% CI]) | 1.8 [1.2, 2.4] | 5.3 [4.3, 6.3] | 12.1 [10.5, 13.7] | 18.5 [14.6, 22.4] | $F_{(3,132)} = 92.3, p < .001$ | 1<2<3<4 |
| Principal Component of SIPS (Mean [95% CI]) | -1.9 [-2.1, -1.7] | -0.7 [-1.0, -.4] | 2.1 [1.5, 2.7] | 3.7 [2.5, 4.9] | $F_{(3,132)} = 104.1, p < .001$ | 1<2<3<4 |
| Sum of BPRS<sup>†</sup> (Mean [95% CI]) | 14.4 [13.5, 15.3] | 21.2 [19.6, 22.8] | 26.4 [22.9, 29.9] | 21.2 [18.4, 24.0] | $F_{(3,127)} = 28.4, p < .001$ | 1<2~4<3 |
| Principal Component of BPRS (Mean [95% CI]) | -1.3 [-1.5, -1.1] | 0.4 [0, 0.8] | 1.5 [0.6, 2.4] | 0.2 [-0.6, 1.0] | $F_{(3,127)} = 26.3, p < .001$ | 1<2~4<3 |

<sup>†</sup> Excluding items related to psychotic disorders (Hafkenscheid, 1991)
*Note:* ADHD = Attention Deficit and Hyperactivity Disorder, GAF = Global Assessment of Functioning
*Note.* Group differences in age and gender were statistically significant and were therefore included as covariates in follow-up analyses. No other demographic variables differed significantly between groups.

To reduce potential overlap with HC, participants who met criteria for a psychiatric diagnosis yet were not help-seeking (*N* = 6) were excluded from categorical analyses but retained for dimensional analyses. Similarly, to reduce potential overlap with psychosis-vulnerability, participants who met criteria for schizotypal personality disorder but not for CHR-P criteria (*N* = 10) were excluded from categorical analyses but retained for dimensional analyses.

Recruitment sources included university students, community advertisements, referrals to the early-identification clinic, the psychiatric division at Rambam Medical Center, and a community mental-health NGO (see Table S1 for distribution of recruitment sites).

#### 2.3.3 Psychosis-spectrum symptoms

To obtain a single index of psychosis-spectrum symptom severity, we conducted a principal component analysis (PCA) on SIPS positive-symptom scores and summary statistics that account for their non-ordinal scaling. The PCA yielded a single psychosis-spectrum component (∼70% variance explained; Fig. S1A).

#### 2.3.4 General psychiatric symptoms

General psychopathology independent of psychotic symptoms was assessed using a composite BPRS score excluding psychosis-related items (Hafkenscheid, 1991). Principal component analysis yielded a single BPRS component (∼90% variance explained; Fig. S1B).

### 2.4 Experimental Paradigm and Cognitive / Metacognitive Measures

#### 2.4.1 Experimental procedure

Participants completed a probabilistic learning task (Harrison et al., 2021; Iglesias et al., 2013) adapted to VR and described in detail in Stern et al. (2026). On each of 160 trials, participants viewed one of two colored squares that, according to an implicit probabilistic color-location rule, indicated the more probable target location (left/right). Participants explicitly predicted the target location and entered the virtual environment, where the target (butterfly) appeared after 300 milliseconds. Gaze analyses focused exclusively on this 300 milliseconds interval (Fig. 1A).

**Fig. 1.**
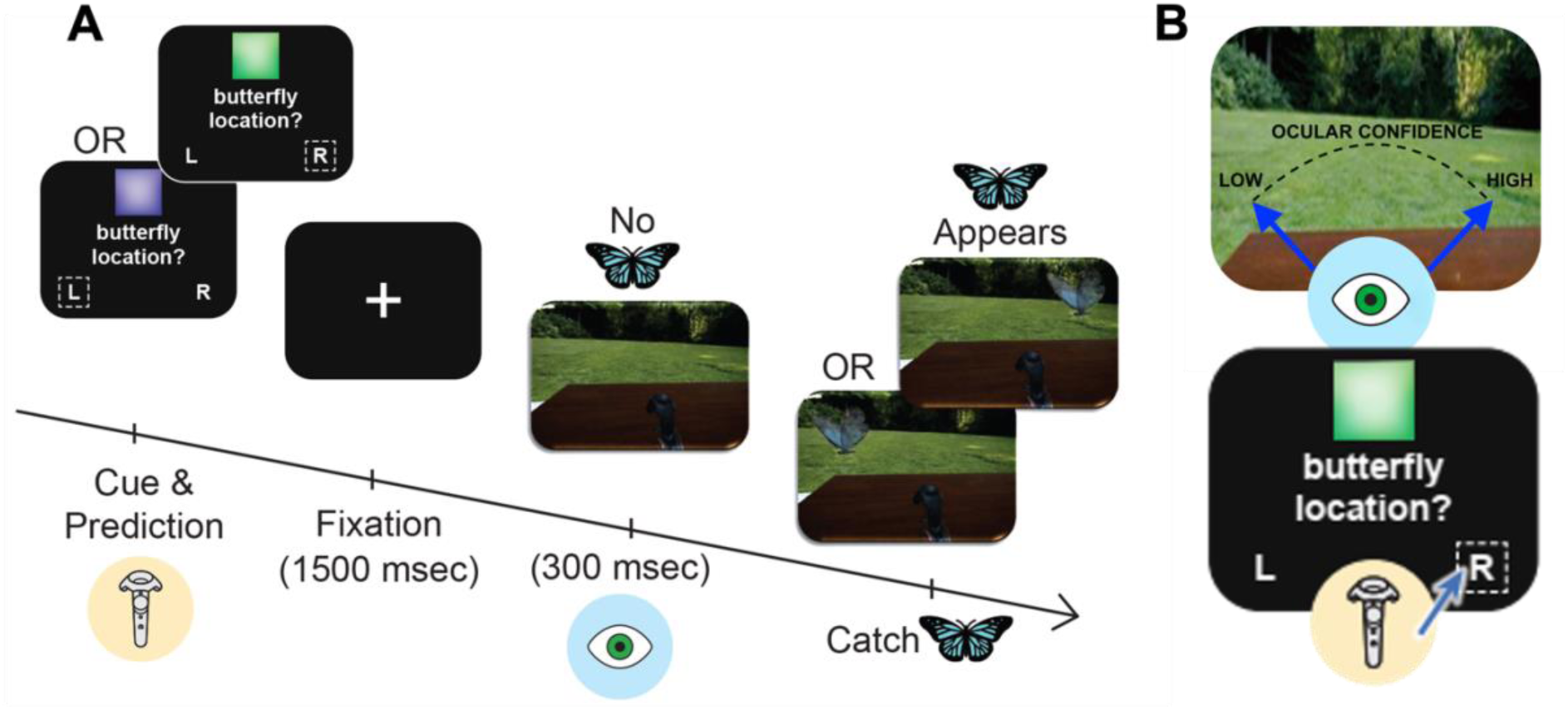
Experimental paradigm and gaze-based confidence measures. (A) Trial structure of the virtual reality–based probabilistic learning task. On each trial, participants predicted the upcoming target location (left/right) using the handheld controller. After a fixation period, they entered the virtual environment and after an interval the target appeared. Post-decision gaze direction on the horizontal axis was assessed during the 300-millisecond interval preceding target appearance. (B) Gaze-based confidence, defined as the alignment between the predicted target location and post-decision gaze direction on the horizontal axis. Positive values indicate gaze aligned with the prediction, whereas negative values indicate misalignment.

Participants learned the current color-location mapping rule and used it to predict the upcoming target location. They were informed that the color cues were contingent: if one color predicted the right location, the other predicted the left. Target location followed the rule on 75% of trials and violated it on the remaining 25%. and the rule could change. The rule changed five times across six 32-trial blocks (see SM for details).

#### 2.4.2 Cognitive and metacognitive measures

##### 2.4.2.1 First-order learning

Prediction accuracy was defined as the percentage of trials in which the prediction matched the most probable target location indicated by the underlying rule. Learning strategy was assessed by the proportion of rule switches across consecutive trials (Behrens et al., 2007; Cohen et al., 2007): (1) Lose-Switch, switching after an incorrect prediction (adaptive exploitation), and (2) Win-Switch, switching after a correct prediction (exploration).

##### 2.4.2.2 Implicit metacognitive measures: Gaze-based confidence

Metacognitive performance was indexed by gaze-based confidence, defined as the trial-by-trial alignment between the explicit left-right prediction and the time-weighted direction of post-decision gaze on the horizontal axis during the 300-ms interval preceding target onset (Fig. 1B; see SM for details). Two complementary metacognitive indices were derived from gaze-based confidence, reflecting distinct aspects of metacognitive monitoring: (1) *Overall confidence*, calculated as the mean gaze-based confidence across trials. Higher values indicate greater alignment between post-decision gaze and the predicted target location (higher gaze-based confidence), whereas lower values indicate greater misalignment (lower gaze-based confidence). Together with first-order performance, this measure indexed *confidence calibration*. (2) *Metacognitive sensitivity*, calculated as the difference in gaze-based confidence between correct and incorrect trials (Δ confidence). Positive values indicate greater gaze-based confidence for correct than incorrect decisions.

### 2.5 Statistical Analyses

Statistical analyses were conducted using *R* (version 2023.09.1). Statistical significance was set at *p* < .05 (two-tailed), and assumptions were evaluated prior to analysis. Group differences in demographic and clinical variables were examined using ANOVA, Kruskal-Wallis, or χ² tests, as appropriate. Group differences in H1, H2 and H3a were examined using a one-way ANOVA with Clinical Group (HC, HSC, CHR-P, FEP) as the between-subject factor and the relevant experimental variable was the dependent variable. For the examination of group differences in metacognitive variables (H2 and H3a), we also included accuracy as a covariate to control for differences in first-order performance. For planned pairwise comparisons, both uncorrected *p*-values and Tukey-adjusted *p*-values (adjusted across all pairwise comparisons within the model) are reported for transparency. To quantify evidence for observed null effects (Jeffreys, 1998), Bayesian analyses were performed using the *Bayes-Factor* package (Morey and Rouder, 2023). Dimensional associations between experimental measures and psychosis-spectrum symptom severity were examined using Pearson correlations across all participants. To examine whether metacognitive variables distinguished CHR-P from FEP beyond first-order performance (H3B), we fit nested logistic regression models predicting group membership. Model fit was compared using likelihood-ratio tests.

## 3. Results

### 3.1. Demographic and clinical characteristics of groups

Table 1 presents the demographic and clinical variables. The groups were well-matched on variables such as education and previous exposure to VR. However, the groups differed significantly in age and gender. Consequently, age and gender were included as covariates in subsequent analyses.

As expected, the groups differed significantly in psychosis risk, indexed by both the mean SIPS score and the SIPS principal component. General psychopathology excluding psychotic symptoms, assessed via the BPRS, also differed significantly between groups (see Table S2 and S3 for further details on diagnoses and medications). As expected, general psychopathology was lowest for the HC group relative to the clinical groups. Finally, CHR-P exhibited significantly higher non-psychotic psychiatric symptoms than both HSC and FEP, which did not differ from one another.

### 3.2. Hypotheses Testing

#### 3.2.1 H1: Group differences in learning accuracy

A one-way ANOVA showed a significant effect of Clinical Group on learning accuracy, (*F_3,140_* = 3.12, *p* = .03, η^2^ = 0.06; see Table 2 for summary statistics by group). This effect remained significant after controlling for age and gender via an ANCOVA (*F_3,138_* = 2.8, *p* = .04, η^2^ = 0.06). Accuracy was highest in the HC group and numerically reduced across the clinical groups (see Fig. 2A). Consistent with H1, planned comparisons of HC versus CHR-P and HC versus FEP were both statistically significant, however, only the difference between HC versus FEP remained significant following an adjustment for multiple comparisons (HC vs. CHR-P: *p* = .03; *p _Tukey-adjusted_ = .13;* HC vs. FEP: *p* = .009, *p _Tukey-adjusted_ = .04*; see Table S4 for full results of pairwise comparisons). Across participants, lower accuracy was associated with greater psychosis-spectrum symptom severity (*r* = -.17, *p* = .03; Fig. 2B). To assess specificity to psychosis-spectrum symptoms, SIPS scores were residualized against BPRS scores. The negative association remained significant when using residualized psychosis-spectrum scores (*r* = -.18, *p* = .03; see Fig. S2), indicating that it was not accounted for by shared variance with general psychopathology.

**Table 2.** Learning performance and implicit metacognitive measures by clinical group.

| Variable<br>(Mean [95% CI]) | HC | HSC | CHR-P | FEP |
| --- | --- | --- | --- | --- |
| Accuracy | 67.47<br>[64.86, 70.09] | 65.72<br>[63.12, 68.32] | 62.66<br>[59.09, 66.23] | 60.95<br>[56.01, 65.89] |
| % Win-Switch | 18.59<br>[14.97, 22.21] | 21.75<br>[17.88, 25.63] | 26.64<br>[20.87, 32.40] | 27.68 [19.75, 35.60] |
| % Lose-Switch | 44.06<br>[40.06, 48.06] | 48.30<br>[44.44, 52.16] | 46.38<br>[41.69, 51.07] | 44.90<br>[38.48, 51.32] |
| Overall<br>gaze-based confidence | 0.40<br>[0.35, 0.45] | 0.31<br>[0.26, 0.364] | 0.28<br>[0.205,<br>0.349] | 0.31<br>[0.208, 0.414] |
| Metacognitive<br>Sensitivity | 0.16<br>[0.10, 0.23] | 0.16<br>[0.10, 0.225] | 0.20<br>[0.13, 0.28] | 0.06<br>[−0.04, 0.16] |

**Fig. 2.**
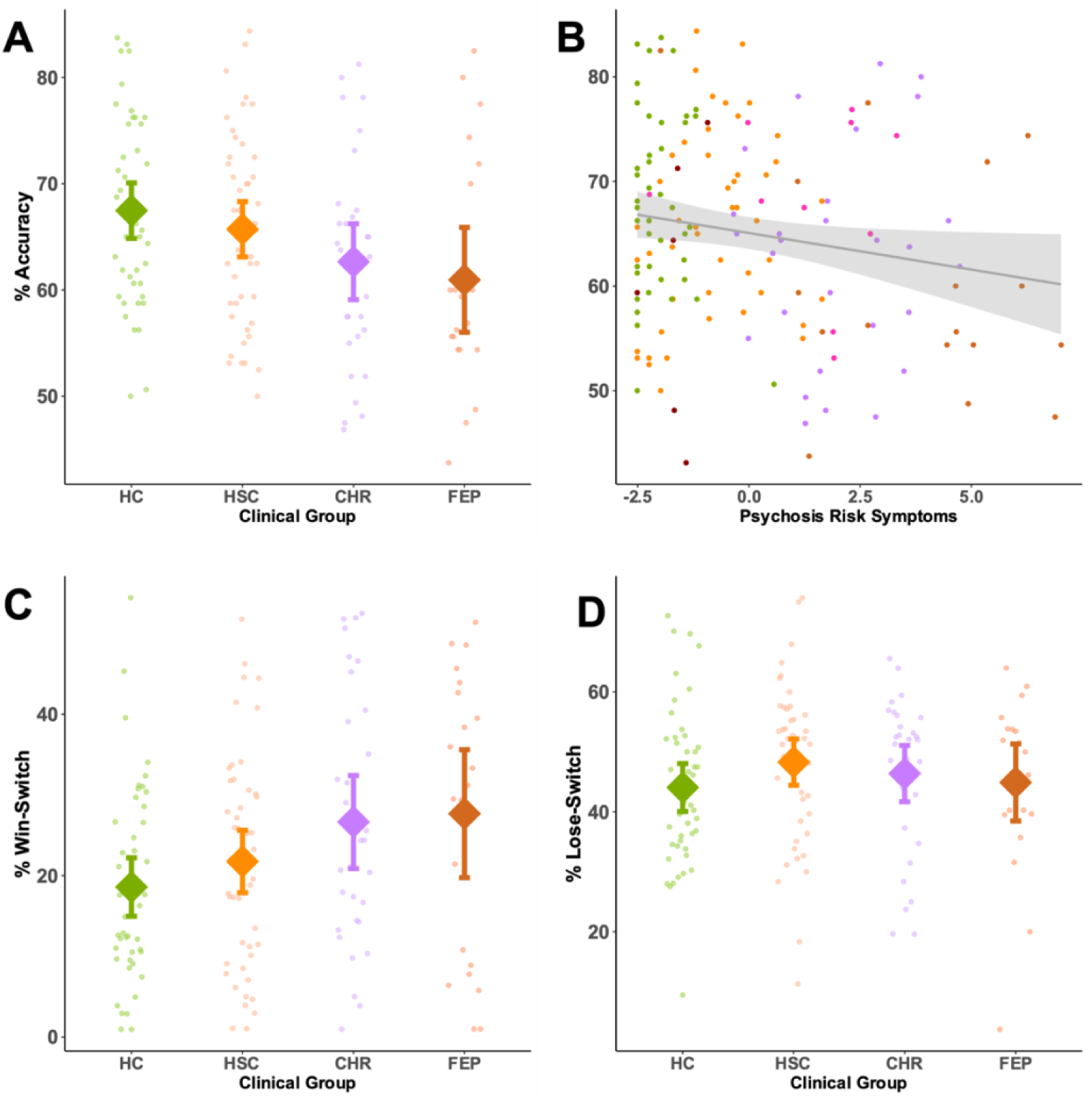
Learning accuracy and exploratory strategies across clinical groups. (A) Prediction accuracy by clinical group. (B) Dimensional association between prediction accuracy and psychosis-spectrum symptom severity across participants. (C–D) Probability of switching the rule underlying one’s prediction following correct (Win-Switch) and incorrect (Lose-Switch) outcomes, by clinical group. Diamonds indicate group means; error bars represent 95% confidence intervals, and dots represent individual participants.

We next compared Win-Switch and Lose-Switch learning strategies. A mixed ANOVA on the proportion of rule switches, with Previous Trial’s Accuracy (Correct / Incorrect) as a within-subject factor and Clinical Group as a between-subject factor revealed a significant interaction, (*F_3,140_* = 3.18, *p* = .03, η^2^ = 0.02, see SM for full results of ANOVA). Planned comparisons showed higher Win-Switch rates in CHR-P and FEP than HC, although neither survived correction for multiple comparisons (HC vs. CHR-P: *p* = .016, *p _Tukey-adjusted_* = .074; HC vs. FEP: *p* = .016, *p _Tukey-adjusted_* = .075; see Table S5). Across participants, greater Win-Switch use was significantly associated with greater psychosis-spectrum symptom severity (*r* = .22, *p* = .007; see Fig S2A). In contrast, the groups did not differ in their use of Lose-Switch (*F*₃,₁₄₀ < 1, *p* = .47), with moderate Bayesian evidence for the absence of group differences (BF₀₁ = 9.19). Across participants, Lose-Switch behavior was not associated with psychosis-spectrum symptoms with moderate Bayesian evidence for the lack of an association (*r* = -.02, *p* = .79, BF₀₁ = 5.14; see Fig S2). Finally, ADHD diagnosis did not account for group differences in learning (see SM).

#### 3.2.2 H2: Group differences in gaze-based confidence calibration

A one-way ANOVA showed a significant effect of Clinical Group on overall confidence, (*F_3, 139_* = 3.16, *p* = .03, η^2^ = 0.06, see Fig. 3A). This effect remained significant after controlling for age and gender (*F*_3, 137_ = 2.70, *p* = .048, η^2^ = 0.06). Overall confidence was highest for HC and reduced for the clinical groups (see Table 2). Consistent with our hypothesis, planned comparisons found a significant reduction for CHR-P relative to HC (*p _uncorrected_* = .005, *p _Tukey-adjusted_* = .03), whereas FEP did not differ significantly compared to HC (*p _uncorrected_* = .07, *p _Tukey-adjusted_* = .27).

**Fig. 3.**
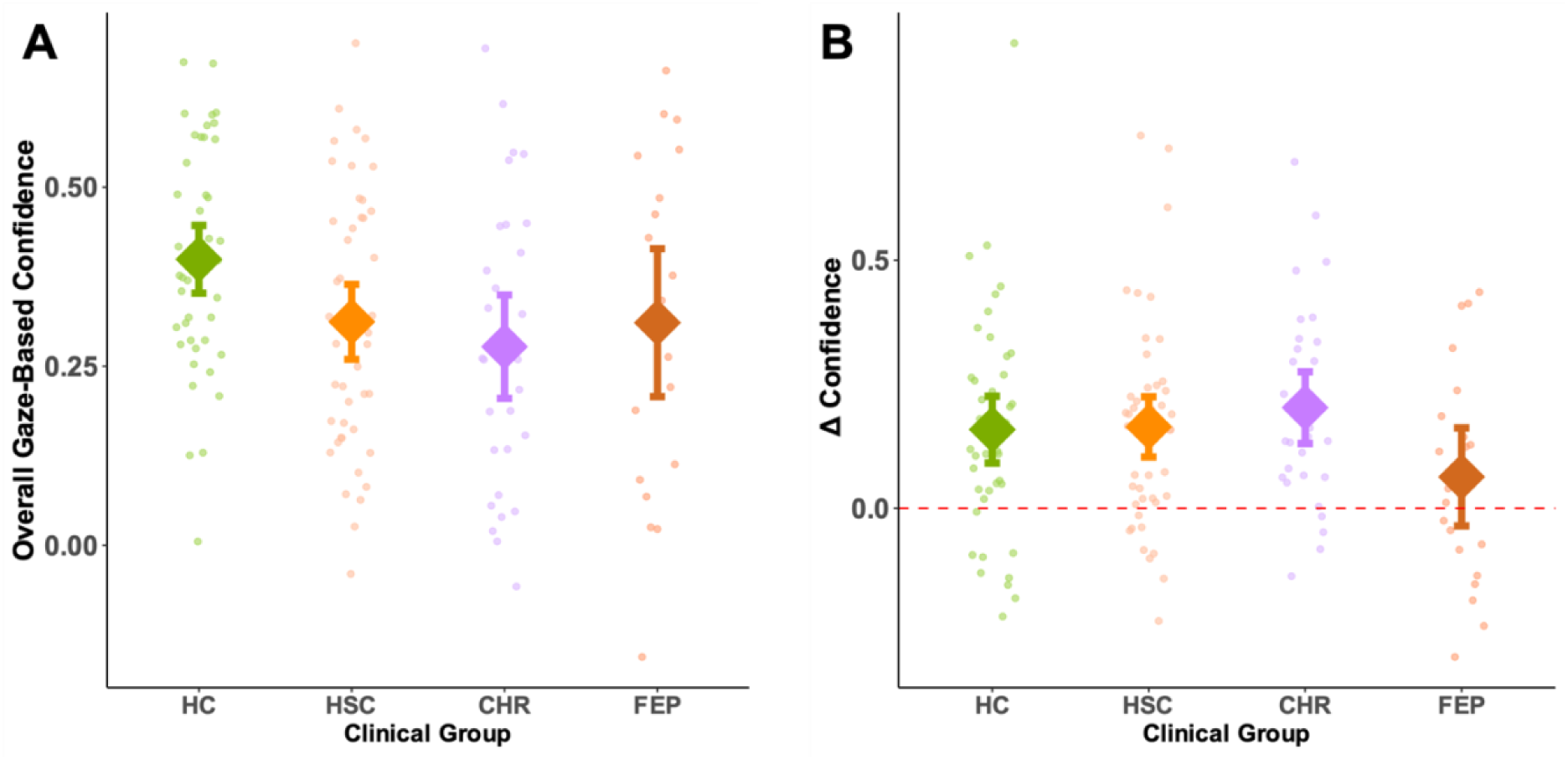
Gaze-based confidence and metacognitive sensitivity across clinical groups. (A) Overall gaze-based confidence averaged across trials by clinical group. Positive values indicate alignment between predictions and post-decision gaze. (B) Metacognitive sensitivity, defined as the difference in gaze-based confidence between correct and incorrect trials, by clinical group. Diamonds represent group mean, error bars represent 95% confidence intervals, and dots are individual participants.

To assess confidence calibration, we repeated the analysis while controlling for first-order accuracy. The overall group effect was no longer significant after accounting for performance and Bayesian analysis provided inconclusive evidence for this lack of difference (*F*₃,₁₃₈ = 1.98, *p* = .12, BF_01_ = 2.20). This pattern was consistent with confidence levels being broadly commensurate with learning accuracy. Consistent with our hypothesis, planned comparisons of residual confidence (confidence adjusted for accuracy) indicated that CHR-P continued to show lower confidence than HC (*p _uncorrected_* = .04), although this did not survive correction for multiple comparisons (*p _Tukey-adjusted_* = .18). In contrast, the FEP group did not differ significantly from HC in residual confidence (*p _uncorrected_* = .10*, p _Tukey-adjusted_* = .47).

#### 3.2.3 H3a: Group differences in gaze-based metacognitive sensitivity

A one-way ANOVA did not show a significant main effect of Group on metacognitive sensitivity (Δ confidence for correct minus incorrect trials; see Methods) with moderate evidence for a lack of a difference (*F_3,139_* = 1.89, *p* = .13, BF_01_ = 3.38, see Fig. 3B). However, inspection of group means suggested a qualitatively distinct pattern across the psychosis continuum. The HC, HSC, and CHR-P groups all exhibited positive metacognitive sensitivity that was significantly greater than zero (all *p’*s < .001), indicating preserved ability to differentiate between correct and incorrect decisions. In contrast, metacognitive sensitivity in the FEP group did not differ significantly from zero (*t*₂₀ = 1.34, *p* = .20). However, Bayesian evidence was inconclusive (BF_01_ = 2.02).

Consistent with this pattern, the HC, HSC, and CHR-P groups did not differ significantly from one another (*F*₂,₁₁₉ < 1.0, *p* = .64), with moderate Bayesian evidence for the absence of group differences (BF₀₁ = 8.70). By contrast, a planned comparison revealed significantly lower metacognitive sensitivity in FEP relative to CHR-P (*t*₄₀,₁₉ = 2.36, *p* = .02). Importantly, this difference remained significant after controlling for first-order learning accuracy (*F*₁,₄₈ = 5.33, *p* = .03), indicating that reduced metacognitive sensitivity in FEP could not be explained solely by impaired task performance.

##### H3b: Implicit gaze-based metacognitive sensitivity adds incremental predictive value for CHR-P versus FEP group membership

Three logistic regression models examined whether metacognitive sensitivity distinguished CHR-P from FEP beyond first-order learning accuracy. Accuracy alone did not predict group membership (OR = 1.19, 95% CI [0.68, 2.15], *p* = .55). In contrast, metacognitive sensitivity alone significantly predicted group status (OR = 2.12, 95% CI [1.14, 4.39], p = .03), with higher sensitivity associated with increased odds of belonging to the CHR-P group. A combined model including both predictors indicated that metacognitive sensitivity remained a significant predictor (OR = 2.16, 95% CI [1.15, 4.55], *p* = .03), whereas accuracy did not (OR = 1.26, 95% CI [0.70, 2.32], *p* = .45). Critically, adding metacognitive sensitivity to the accuracy-only model significantly improved model fit (likelihood-ratio χ²(1) = 5.95, *p* = .015), whereas adding accuracy to the metacognitive-sensitivity model did not (χ²(1) = 0.57, *p* = .45).

## 4. Discussion

### 4.1 Main findings

Consistent with our hypotheses, the present findings reveal a dissociation between first-order learning and implicit metacognition across the psychosis continuum. Although both CHR-P and FEP exhibited impaired probabilistic learning, only FEP exhibited impaired implicit metacognitive calibration and sensitivity. Moreover, implicit metacognitive sensitivity distinguished CHR-P from FEP beyond first-order learning performance. Together, these findings suggest that preserved implicit metacognition may characterize psychosis risk despite impaired learning, whereas disruption of implicit metacognitive monitoring may emerge with the onset of psychosis.

### 4.2 Impaired probabilistic learning across the psychosis continuum

Consistent with previous works in full-blown psychosis (Barch et al., 2017; Pratt et al., 2021) and CHR-P (Cole et al., 2020; Karcher et al., 2019; Millman et al., 2020; Strauss et al., 2021), learning accuracy was reduced in FEP and, to a lesser extent, CHR-P. Importantly, learning impairment was selectively associated with psychosis-spectrum symptoms rather than general psychopathology, and this association remained after controlling for general psychiatric symptoms. Although the sample was diagnostically heterogeneous and ADHD was relatively prevalent, supplementary analyses suggested that these factors did not account for the observed effects, providing preliminary support for the specificity of probabilistic learning impairments to psychosis-spectrum processes.

A leading computational account attributes these learning impairments to overestimation of environmental volatility, resulting in excessive updating in response to ambiguous evidence (Cole et al., 2020; Katthagen et al., 2022). Consistent with this account, higher psychosis-spectrum symptoms were associated with greater reliance on the exploratory Win-Switch strategy, indicating an increased tendency to abandon a correct rule following ambiguous feedback. Such atypical volatility estimation has been proposed to reflect disrupted integration of prior beliefs with sensory evidence, a mechanism implicated in delusion formation (Powers et al., 2025). Although the present cross-sectional design precludes causal inference, these findings suggest that altered volatility estimation may already be present during the psychosis-risk stage.

### 4.3 Metacognitive calibration is preserved in CHR-P but impaired in FEP

CHR-P but FEP participants showed reduced overall gaze-based confidence relative to healthy controls. Importantly, much of this reduction was explained by their diminished learning accuracy, indicating that confidence was broadly commensurate with objective task performance and therefore largely well calibrated. Although there was some evidence of residual under-confidence, this effect was modest and did not survive correction for multiple comparisons. This pattern is consistent with phenomenological accounts of the at-risk state, which emphasize heightened uncertainty and continued questioning of one’s perceptions in the absence of frank psychotic conviction (Parnas, 2012; Sass and Parnas, 2003). It also contrasts with reports of increased explicit confidence in schizotypy and related traits (Lehmann and Ettinger, 2023; Rouault et al., 2018), suggesting that implicit and explicit measures may capture partially distinct aspects of metacognitive functioning. By contrast, FEP participants did not exhibit a corresponding reduction in gaze-based confidence despite impaired learning. This mismatch between performance and confidence suggests impaired confidence calibration, consistent with longstanding accounts of deteriorating insight and uncertainty monitoring following the onset of psychosis (Aleman et al., 2006; Wright et al., 2023).

### 4.4 Metacognitive sensitivity is preserved in CHR-P but impaired in FEP

Our most important metacognitive finding was that gaze-based metacognitive sensitivity was preserved in CHR-P but impaired in FEP, despite comparable first-order learning deficits in the two groups. This pattern aligns with a central clinical feature of the at-risk state, namely relatively preserved insight. Whereas CHR-P participants retained the ability to distinguish correct from incorrect decisions through their gaze behavior, FEP participants showed markedly reduced metacognitive sensitivity. Importantly, what differentiated CHR-P from FEP was not the presence of learning deficits per se, as both groups exhibited impaired task performance, but rather the ability to recognize when those deficits had resulted in an incorrect decision. These findings are consistent with clinical observations that insight and uncertainty monitoring often remain relatively preserved in CHR-P but deteriorate following psychosis onset.

The finding that metacognitive sensitivity distinguished CHR-P from FEP beyond first-order learning further suggests that second-order monitoring provides clinically relevant information that is not captured by cognitive performance measures alone. However, because the present study is cross-sectional, the temporal interpretation of these findings remains uncertain. Longitudinal studies will be essential to determine whether diminished metacognitive sensitivity precedes the transition to psychosis or emerges as a consequence of illness onset.

An alternative interpretation is that reduced gaze-based metacognitive sensitivity reflects a broader decoupling between gaze behavior and task-relevant information rather than impaired metacognitive monitoring. This possibility warrants consideration because abnormalities in gaze control and attentional allocation have been documented across the psychosis spectrum (Benson et al., 2012; Miura et al., 2025; Thakkar et al., 2017). However, such an account would predict broadly reduced gaze–prediction coupling, particularly in FEP. Instead, FEP participants exhibited overall gaze-based confidence comparable to healthy controls but selectively impaired metacognitive sensitivity. This pattern is therefore more consistent with disrupted uncertainty monitoring than with a generalized oculomotor or attentional abnormality. Future studies combining gaze-based confidence with explicit confidence ratings and independent measures of gaze control will be important for further disentangling metacognitive and sensorimotor contributions.

### 4.5 Strengths and Limitations

A major strength of the present study is the use of a novel implicit behavioral measure of metacognition based on post-decision gaze. Unlike explicit confidence ratings, gaze-based confidence captures uncertainty monitoring without requiring introspection or interrupting task performance, providing an ecologically grounded measure of metacognitive functioning.

Several limitations warrant consideration. First, although the overall sample was relatively large, the CHR-P and FEP subgroups were modest in size, and the FEP group was diagnostically heterogeneous, warranting caution when generalizing these findings. Second, the cross-sectional design precludes conclusions regarding the temporal relationship between metacognitive alterations and illness progression. Finally, explicit confidence ratings were not obtained in the clinical groups, precluding direct comparison between implicit and explicit metacognitive measures. Future longitudinal studies combining both measures will be important for clarifying how different aspects of metacognition evolve across the psychosis continuum.

## 5. Conclusions

The present findings suggest a dissociation between first-order learning and implicit metacognitive monitoring across the psychosis continuum. Preserved implicit metacognitive functioning may characterize psychosis risk, whereas its disruption may accompany psychosis onset. Implicit behavioral measures such as gaze-based confidence may therefore refine clinical characterization. Longitudinal studies are needed to determine their prognostic and therapeutic value.

## Author contributions

**Yonatan Stern**: Conceptualization, Data curation, Formal analysis, Investigation, Methodology, Project administration, Resources, Software, Validation, Visualization, Writing – original draft, Writing – review and editing. **Donia Sussan**: Investigation, Methodology, Project administration. **Barnaby Nelson**: Conceptualization, Methodology, Writing – review and editing. **Uri Hertz**: Conceptualization, Methodology, Writing – review and editing. **Morris Goldsmith**: Conceptualization, Methodology, Writing – review and editing. **Eyal Bergmann**: Investigation, Project administration, Writing – review and editing. **Lauren Nashashibi**: Investigation, Project administration, Writing – review and editing. **Danny Koren**: Conceptualization, Data curation, Methodology, Project administration, Resources, Software, Funding Acquisition, Supervision, Writing – original draft, Writing – review and editing. **Roy Salomon**: Conceptualization, Data curation, Methodology, Resources, Software, Funding Acquisition, Supervision, Writing – original draft, Writing – review and editing.

## Declaration of generative AI and AI-assisted technologies in the manuscript preparation process

During the preparation of this work the authors used ChatGPT and Gemini in order to assist in language editing. After using these tools, the authors reviewed and edited the content as needed and take full responsibility for the content of the published article.

## Acknowledgments

We express our gratitude to the patients and their families for their participation. We also thank Geffen Markusfield and Annael Halimi for their help with administration of the clinical interviews, and Natali Manov, Shani Berkovitch, Neta Chernikov, Shoham Buchnik, Eiman Egbaria members of the ’Bridge Over Troubled Waters’ lab for their help with administering the experimental tasks, as well as the staff of Rambam’s Psychiatry Division for their support in the recruitment process.

