## Supplemental Material for "Preserved implicit metacognitive sensitivity distinguishes psychosis risk from first-episode psychosis: A cross-sectional virtual reality-based study"

**Methods**

**Clinical Assessments**

In addition to the measures described in the main text, participants were also administered an abbreviated version of the Examination of Anomalous Self-Experiences (EASE;(Parnas et al., 2005)) to evaluate Basic Self-Disturbances (BSD). Informed by theoretical phenomenological considerations, clinical expertise, and empirical findings, we utilized a 10-item subset (EASE-10) designed to capture the most prototypical disturbances of the basic self. This version was previously implemented (Koren et al., 2013), received conceptual validation from co-developer Josef Parnas, and maintains a high correlation with the comprehensive interview format. The specific items assessed were: hyperreflexivity, loss of common sense/perplexity, mirror-related phenomena, loss of thought ipseity, spatialization of experience, ambivalence, diminished sense of basic self, loss of first-person perspective, perceptualization of inner speech, and thought pressure. The item assessing ‘motor disturbances’ was also included for exploratory purposes.

In addition to the clinical interviews, participants completed a battery of self-report questionnaires. These included the Prodromal Questionnaire–Brief (PQ-B) to screen for attenuated psychotic symptoms (Loewy et al., 2011), and the Inventory of Psychotic-Like Anomalous Self-Experiences (IPASE) to evaluate disturbances in self-experience (Cicero et al., 2017). To account for comorbid symptoms and history, we also administered the Obsessive-Compulsive Inventory–Revised (OCI-R) (Foa et al., 2002), the Adverse Childhood Experiences (ACE) questionnaire (Felitti et al., 1998), the DSM-5 Level 1 Cross-Cutting Symptom Measure for general psychopathology (Clarke and Kuhl, 2014), and the Alcohol, Smoking and Substance Involvement Screening Test (ASSIST) to monitor substance use (Group, 2002).

**Recruitment Sites**

Table S1 provides a breakdown of recruitment sources for each clinical group, distinguishing between participants recruited via community-based frameworks and those referred through the Psychiatric Division at Rambam Medical Center. Community frameworks are defined here as non-clinical recruitment channels, which include university student cohorts, public community advertisements, and outreach through a local mental-health NGO.

| **Recruitment Source** | **Healthy Controls (HC)** | **Help-Seeking Controls (HSC)** | **Clinical High Risk (CHR)** | **First-Episode Psychosis (FEP)** | **Schizotypal Personality Disorder (SPD)** | **Non-help seeking Controls** |
| --- | --- | --- | --- | --- | --- | --- |
| **Community Frameworks** | 45 | 21 | 5 | 3 | 7 | 6 |
| **Psychiatric Division** | 0 | 26 | 26 | 18 | 3 | 0 |

**Table S1.** Participant distribution across recruitment frameworks by clinical group classification.

**Clinical Measures**

**Principal component analysis of psychosis-spectrum symptom severity**

To derive a latent index of symptom severity, a Principal Component Analyses (PCA) were conducted for the SIPS. This approach accounts for the non-linear, qualitative shifts inherent in these scales, where scores above specific thresholds (e.g., >2 for SIPS) represent a clinically significant transition. By entering both the aggregate sum and the total count of above-threshold symptoms into the PCA, the resulting first principal component parsimoniously captures both the overall symptom burden and the qualitative shift toward clinical-level pathology.

The SIPS PCA was derived from the sum score across all subscales, the total count of above-threshold symptoms, the score on each individual subscale, and the number of schizotypal personality disorder symptoms endorsed. Importantly, the first principal component captured over 70% of the variance (see Fig. S1A) and was highly correlated to the aggregate sum (*r* = .99, *p* < .001).

**Principal component analysis of general psychiatric symptoms**

To derive a measure of general psychopathology that is independent and cdistinct from psychosis-spectrum severity, we focused the BPRS items that are not associated with schizophrenia spectrum disorders (i.e., items Hostility, Grandiosity, Suspiciousness, Hallucinatory Behavior, Unusual Thought Content, Conceptual Disorganization, Blunted Affect, Emotional Withdrawal, Uncooperativeness, and Mannerisms and Posturing; Hafkenscheid, 1991). The PCA was derived from the aggregate sum. the total count of above-threshold items, and count of items that were endorsed. Importantly, the first principal component captured over 90% of the variance (see Fig. S1B) and was highly correlated to the aggregate sum (*r* = .99, *p* < .001).

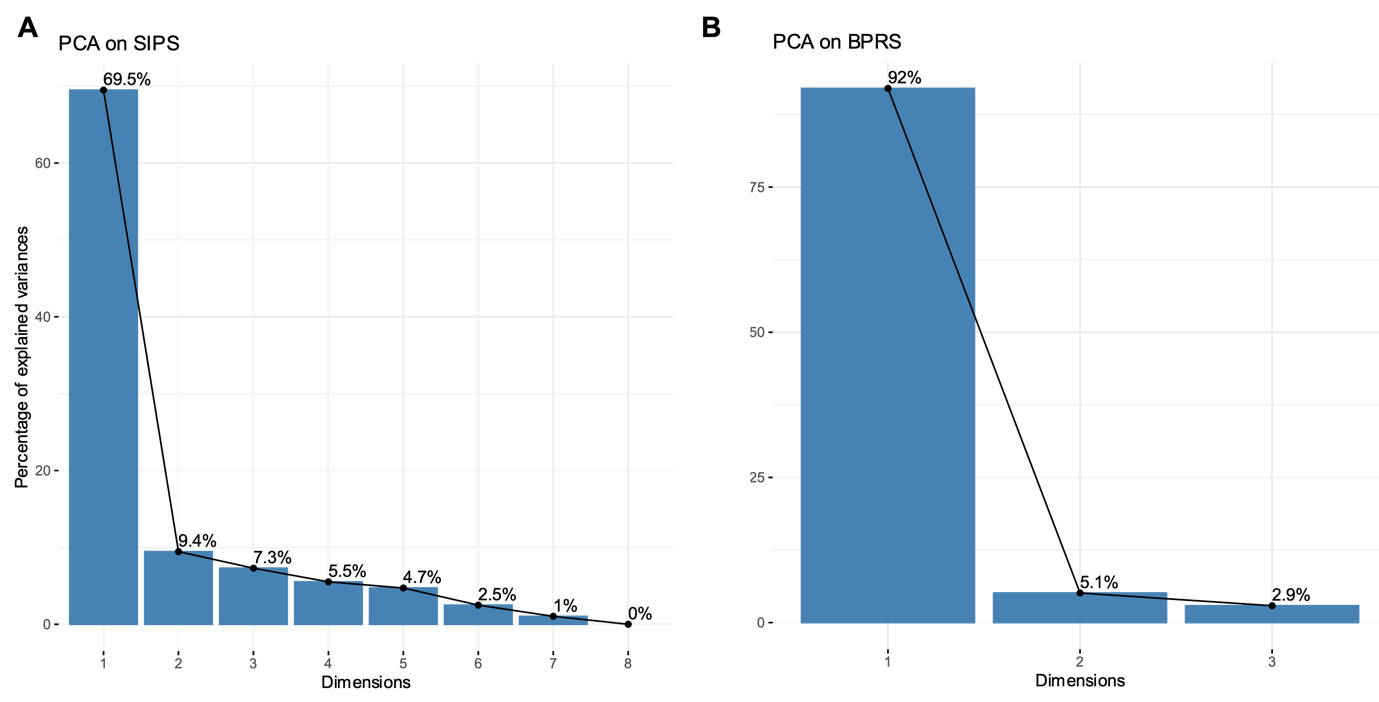

**Figure S1. Scree plots showing the percentage of explained variance for the clinical symptom PCAs.** (A) Principal Component Analysis of SIPS positive symptoms, where the first component accounts for 69.5% of the total variance. (B) Principal Component Analysis of general psychopathology (excluding schizophrenia spectrum associated items), with the first component accounting for 92% of the total variance. For both PCAs the first component explains most of the variance and after it there is a pronounced “elbow”, justifying its use as a parsimonious index of symptom severity.

**Task description**

The experimental procedure employed a well-established probabilistic learning task (Harrison et al., 2021; Iglesias et al., 2013; Stern et al., 2025), adapted to immersive virtual reality (VR) environment. The environment was built using Unity (v2018.3.2), and participants wore a HTC Vive Pro Eye head-mounted display monitor with a 90 Hz refresh rate. The virtual environment was designed to align with the participant’s physical workspace, where they remained seated at a table throughout the experiment.

The session consisted of 160 trials. Each trial began with the presentation of one of two probabilistic spatial cues (colored squares). Participants were required to predict the upcoming target’s location (Right vs. Left) via the handheld controller. Following this prediction and a fixation period of 1.5 seconds, participants transitioned into a virtual environment where the target, an animated butterfly, appeared after a 300 ms delay. The target appeared equally often in one of six possible locations distributed symmetrically across the left and right sides of the environment. The target was static and made animated flapping movements. Participants were instructed to reach toward the target as quickly and accurately as possible once it was identified.

Participants were informed that: (1) an underlying probabilistic rule governed the association between cue color and target location, and the rule was valid on 75% of trials. (2) The cues were contingent and mutually exclusive, such that if Color A signaled a leftward target, Color B necessarily signaled a rightward target under a given rule. (3) The underlying rule could be changed multiple times throughout the session, and they need to detect these changes and respond accordingly. In reality, the rule changed five times during the experiment, with each block consisting of 32 consecutive trials that were governed by the same rule. To maintain experimental consistency, a set of predetermined stimulus sequences was utilized across all participants.

**Gaze-based confidence**

In previous work in the general population, we developed and validated a behavioral manifestation of decision confidence based on post-decision gaze, that we coined Gaze-based confidence (Stern et al., 2025). In brief, the graded alignment between a participant's post-decision gaze direction and their explicit prediction regarding the target’s location (right/left), we derived a continuous measure termed Gaze Prediction. Gaze direction was measured during a preregistered 300 ms interval upon entry into the virtual environment and concluding at the onset of the target stimulus. To account for individual variability, horizontal gaze eccentricity was first normalized relative to the viewer-centered midpoint. We then calculated a time-weighted average of these normalized values for each trial to yield a composite index of the duration and magnitude of lateral gaze bias. Gaze Prediction was derived by adjusting the sign of the gaze direction to indicate convergence with (positive) or divergence from (negative) the predicted target direction. This was achieved using the following algorithm:

$$Gaze Prediction=\left\{ \begin{aligned} G\mathrm{aze}\mathrm{Normalized} if Prediction ="Right" \\ G\mathrm{aze}Normalized*-1 if Prediction ="Left" \end{aligned} \right.$$

Conceptually, this measure indexes the strength of alignment between post-decision gaze and the preceding binary choice. Positive values scale the degree of gaze bias in the predicted direction, while negative values scale the degree of misalignment toward the opposite direction. Importantly, this implicit measure exhibited partial overlap with decision confidence in a number of converging manners. It was modestly correlated to explicit confidence, exhibited the computational hallmarks of confidence (Sanders et al., 2016), and robustly correlated to the model-based decision certainty.

**Results**

### Clinical Characteristics

| **Group** | **Anxiety** | **Depression** | **OCD** | **Other*** | **Personality Disorders** | **Psychotic**  **Disorders** |
| --- | --- | --- | --- | --- | --- | --- |
| **HSC** | 21% (10) | 38% (18) | 17% (8) | 38% (18) | 17% (8) | - |
| **CHR-P** | 42% (13) | 32% (10) | 23% (7) | 26% (8) | 26% (8) | - |
| **FEP** | 10% (2) | 5% (1) | 24% (5) | 24% (5) | 14% (3) | 95% (20) |

*** Bipolar, Eating disorder, Posttraumatic Stress Disorder, Adjustment Disorder, Somatoform Disorder, Tic Disorder.

**Table S2.** Prevalence (and Number of Subjects) of diagnosis by Clinical Group

| Clinical Group | Antidepressants | Mood Stabilizers | Antipsychotics | Sedatives | Other Medications |
| --- | --- | --- | --- | --- | --- |
| HSC | 45% (20) | 18% (8) | 14% (6) | 11% (5) | 5% (2) |
| CHR-P | 39% (11) | 7% (2) | 29% (8) | 18% (5) | 4% (1) |
| FEP | 33% (7) | 10% (2) | 57% (12) | 38% (8) | 19% (4) |

**Table S3**. Psychiatric medication by clinical group

**H1: Group differences in learning accuracy**

| **Clinical Group 1** | **Clinical Group 2** | ***p* _(Unadjusted)_** | ***p _(_* _Tukey-adjusted)_** |
| --- | --- | --- | --- |
| HC | HSC | 0.368 | 0.803 |
| HC | CHR | 0.028 * | 0.125 |
| HC | FEP | 0.009 ** | 0.044 * |
| HSC | CHR | 0.158 | 0.490 |
| HSC | FEP | 0.053 | 0.212 |
| CHR | FEP | 0.517 | 0.916 |

**Table S4.** Pairwise comparisons of mean accuracy between clinical groups.

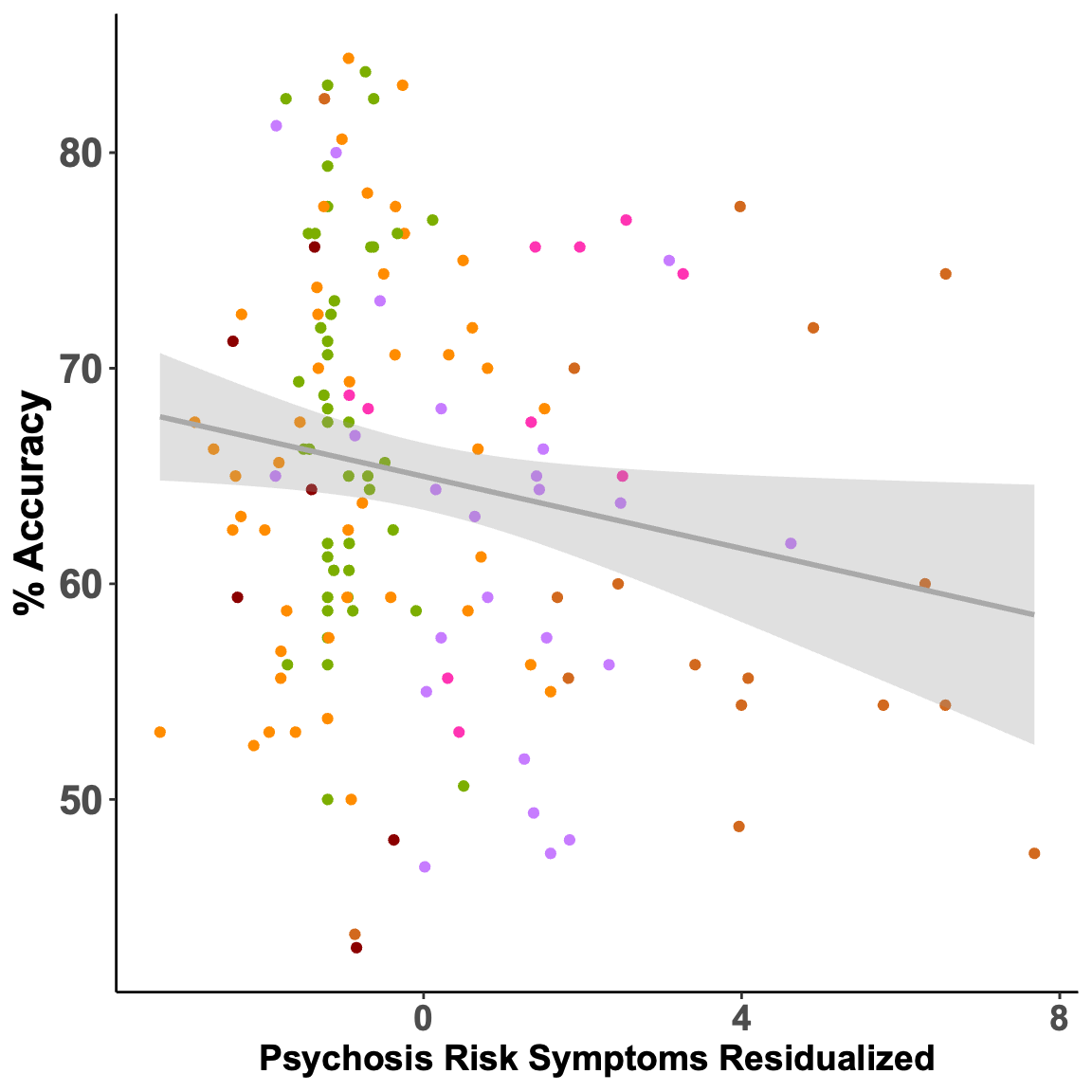

**Fig. S2. Association between learning accuracy and residualized psychosis-spectrum symptoms.** A dimensional analysis across all participants revealed that lower accuracy was significantly associated with greater psychosis-spectrum symptom severity (*r* = -0.18, *p* = 0.03), even when residualizing from the SIPS principal component the BPRS principal component that reflects general psychopathology. Individual data points are colored by clinical group, with the solid grey line representing the linear regression fit and the shaded area indicating the 95% confidence interval.

**Group differences in learning strategies**

As noted in the main text, to investigate the group differences in accuracy, we compared their use of the Win-Switch and Lose-Switch learning strategies. A mixed ANOVA with Previous Trial’s Accuracy (Correct / Incorrect) as a within-subject factor and Clinical Group as a between-subject factor revealed a significant interaction, (F_3,140_ = 3.18, *p* = .03, η^2^ = 0.02). As expected, there was a robust main effect of previous trial’s accuracy on switching such that after errors participants were more likely to switch the rule governing their response (F_1,140_ = 326.58, *p* < .001, η^2^ = 0.38). In contrast the main effect of Clinical Group on overall switching was not significant (F_3,140_ = 1.58, *p* = .20).

| **Clinical Group 1** | **Clinical Group 2** | ***p* _(Unadjusted)_** | ***p _(_* _Tukey-adjusted)_** |
| --- | --- | --- | --- |
| HC | HSC | 0.285 | 0.706 |
| HC | CHR | 0.016 * | 0.074 |
| HC | FEP | 0.016 * | 0.075 |
| HSC | CHR | 0.137 | 0.442 |
| HSC | FEP | 0.112 | 0.382 |
| CHR | FEP | 0.795 | 0.994 |

**Table S5. Pairwise comparisons of Win-Switch trial proportions between clinical groups.** Win-Switch trials are defined as those in which the participant’s previous response was correct, yet they subsequently switched their underlying response rule.

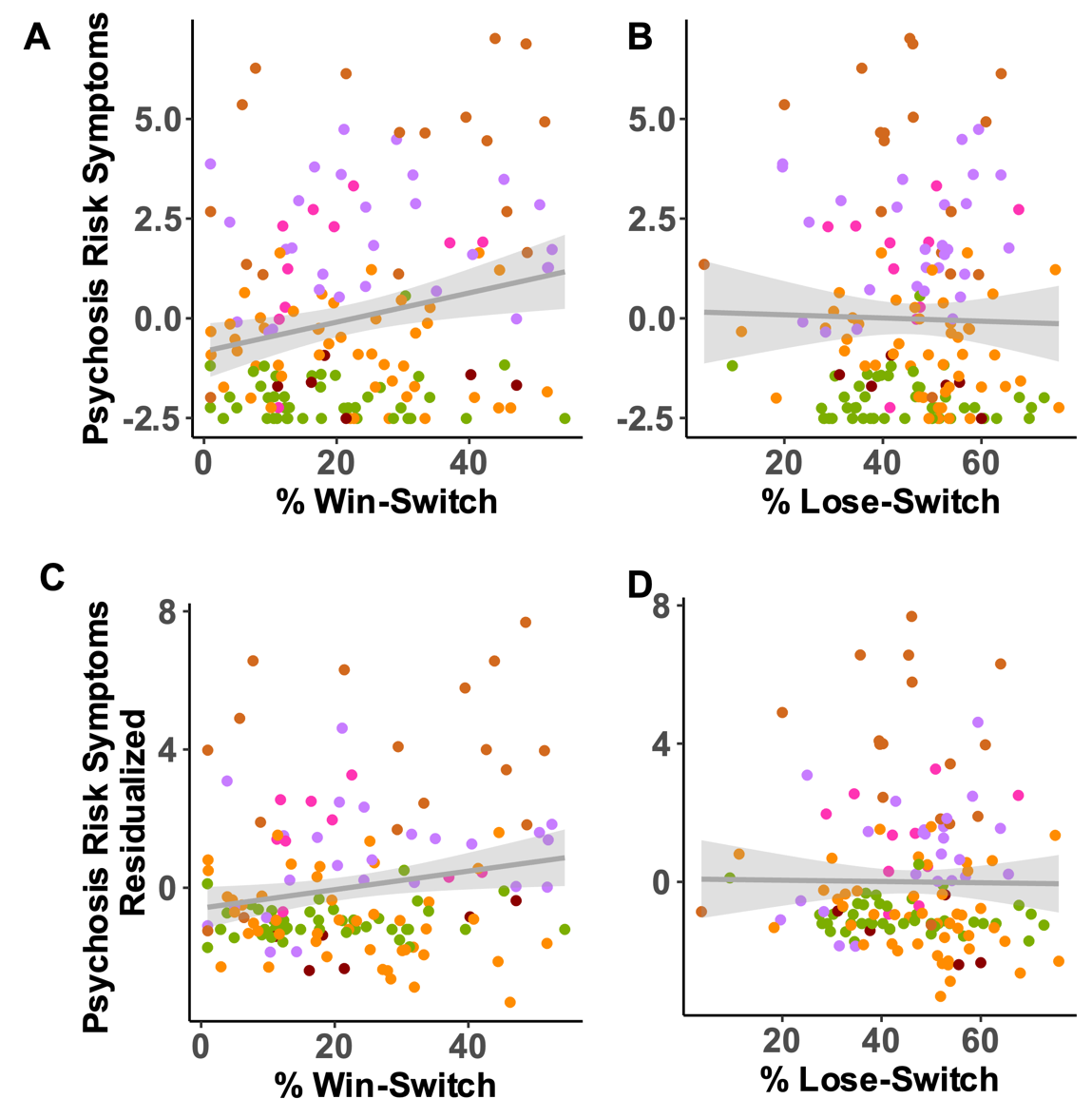

**Figure S2. Dimensional association between learning strategies and psychosis-spectrum symptom severity.** (A) Association between the proportion of Win-Switch trials and psychosis-spectrum symptom severity, revealed a significant positive correlation (*r*= .22, *p* = .007). (B) Association between the proportion of Lose-Switch trials and psychosis-spectrum symptoms did not find a significant association (*r*= -.02, *p* = .79, BF₀₁ = 9.19). Individual data points represent single participants, colored by their respective clinical group. The solid grey line indicates the linear regression fit, and the shaded area represents the 95% confidence interval. (C-D) Examine the correlation with psychosis-spectrum symptoms that are residualized from general psychopathology. The correlation to Win-Switch was significant (*r* = .19, *p* = .02; panel C), whereas the correlation to Lose-Switch was not significant (*r* = -.01, *p* = .89, BF₀₁ = 5.19; panel D).

**Effect of ADHD on learning**

Given the relatively high prevalence of ADHD diagnoses in the sample, we conducted additional analyses to examine its potential confounding effect. Participants with ADHD did not significantly differ in learning accuracy and a Bayesian analysis provided moderate evidence (*t*_90.55_ = -0.14, *p* = .89, BF₀₁ = 5.23). Likewise including ADHD diagnosis as a covariate in the ANOVA examining group differences in accuracy it was not significant and Bayesian analysis provided moderate evidence for a lack of association (F_1,135_ < 1, *p* = .42, BF₀₁ = 4.09). ADHD status was not significantly associated with learning accuracy (*r* = 0.03, *p* = .75), while the main effect of clinical group remained unchanged (F_35,13549_ = 2.353.04, *p* = .034). In addition, a mixed-design ANOVA revealed no evidence for a Task Half × ADHD Status interaction and Bayesian analysis provided moderate evidence for a lack of interaction ((F_1,154_ < 1, *p* = .97, BF₀₁ = 4.52), , suggesting sting that the group differences in accuracy were not driven by performance decay over time.

**Pre-registered Exclusion Criteria and Sample Size**

### Section S1. Exclusion Criteria and Deviation from the Pre-Registration

In our pre-registration ([https://osf.io/bc9np/overview](https://www.google.com/search?q=https://osf.io/bc9np/overview)), we specified several exclusion criteria designed to ensure data quality and reliability that were based on previous work in the general population (Stern et al., 2025). In brief these criteria were:

1. **Overall Accuracy:** Accuracy (explicit response matching target location) below 50% for model-free analyses, or below 55% for model-based analyses.
2. **Validity Effect:** Accuracy on "Valid" trials must be greater than accuracy on "Invalid" trials.
3. **Sufficient Proportion of Valid Trials:** Participants were excluded if they failed to provide a sufficient proportion of valid trials (< 70 % for HC group; < 65 % for the other groups). A trial was considered valid only if it survived all trial-level filters. Specifically, those in which a target was presented, an explicit response was made within the defined reaction time window (250 – 5000 ms and within 3 SD of the participant’s mean reaction time), and the absence of technical VR malfunctions or significant ocular data corruption.
4. **Instructional Compliance:** Exclusion based on experimenter reports of non-compliance prior to data analysis.

A total of 164 participants completed the probabilistic learning task and provided sufficient clinical data for group assignment. Importantly, we deviated from our pre-registerion and did not exclude participants based on the performance criteria (i.e., criteria 1 and 2). These criteria, which were derived from a previous study in healthy participants from the general population (Stern et al., 2025), proved to be confounded with the clinical impairments inherent to the clinical groups. Applying these criteria would have resulted in systematic, skewed exclusions, disproportionately removing the most impaired participants and potentially masking the very deficits under investigatio. In addition, they would significantly reduce the sample size and reduce the statical power of analyses concerning the primary groups of interest (i.e., CHR-P and FEP). Below, we describe the statistical and methodological considerations for this decision.

The application of pre-registered performance criteria (i.e., accuracy threshold and cue-validity) resulted in a significant and non-random reduction of the sample by clinical group (see Table S1). While exclusion was minimal for Healthy Controls (HC; 6.5%) and Help Seeking Controls (HSC; 4.2%), it was substantially higher in the Clinical High Risk for Psychosis (CHR-P; 22.6%) and First Episode Psychosis (FEP; 19.0%) groups.

To formally assess this differential exclusion rate, we conducted a Chi-square test of independence, which revealed a significant association between clinical group and exclusion status χ² = 8.896, p = .031). Given the small cell counts in specific groups, we also calculated Fisher’s Exact Test, which confirmed this significant group difference (*p* = .026). Furthermore, the remaining sample sizes for the primary groups of interest (i.e., CHR-P and FEP) would have been insufficient to maintain adequate statistical power or provide a representative characterization of these populations. Consequently, we did not utilize these criteria.

| Clinical Group | Initial *N* | Exclusion  Criteria 1 | Exclusion  Criteria 2 | Total Excluded | Final *N* | % of Group Lost |
| --- | --- | --- | --- | --- | --- | --- |
| HC | 46 | 3 | 0 | 3 | 43 | 6.5 |
| CC | 48 | 2 | 0 | 2 | 46 | 4.2 |
| CHR | 31 | 4 | 3 | 7 | 24 | 22.6 |
| FEP | 21 | 4 | 1 | 4 | 17 | 19.0 |

**Table S6.** Participant attrition and analytic sample determination across groups when applying pre-registered inclusion criteria of learning performance.

Following the third criterion, two participants were removed: one participant from the CHR-P group who had an insufficient number of valid, and one participant from the HC group had a corrupted behavioral output file. Additionally, one participant from the HC group provided an insufficient number of trials with valid ocular data and was excluded from ocular analyses, while their explicit behavioral responses were retained. Following the fourth criterion, two participants ( one from HC and one from HSC) were excluded based on experimenter reports of non-compliance with task instructions during the session. Consequently, the final sample for the primary behavioral analysis consisted of 160 participants.

Furthermore, we slightly deviated from the pre-registered sample size requirements. For the CHR-P group, we included 30 participants, exceeding the pre-registered minimum of 25, to ensure alignment with a parallel sense of agency study ([https://osf.io/ubjeq](https://www.google.com/search?q=https://osf.io/ubjeq)) that this cohort also completed. Finally, as outlined in our current pre-registration, we included the FEP group (*N* = 20) due to its theoretical significance, having reached the pre-registered minimum requirement.

**Results of Pre-registered Analyses**

Our pre-registered hypotheses were initially informed by exploratory findings from a non-clinical cohort. However as reported below, these patterns did not replicate within the present clinical population. Following a preliminary analysis of the current clinical data and a comprehensive review of the literature concerning psychosis-risk, we shifted the primary focus of the main manuscript to better align with the established findings and specific characteristics of clinical groups. Nevertheless, for the sake of transparency, we report the results of the original pre-registered analyses in this section.

### 3.2.1 H1: Group differences in eye-motor switching

### Our primary hypothesis (H1) posited that "across the psychosis continuum, ocular learning entails increased maladaptive motor perseverance." This was evaluated using two distinct measures. First, H1a employed a model-free ocular switching rate, defined as the proportion of trials in which gaze was directed toward a different location (i.e. right/ left) than on the preceding trial. Second, H1b examined ocular perseverance using a model-based parameter (⍴).

### 3.2.1 H1a: Group differences in model-free ocular motor switching rate

H1a predicted a reduced rate of ocular motor switching in the CHR-P group relative to HC, and a significant relationship between lower switching rates and increased psychosis-proneness symptoms. Contrary to our hypothesis, a one-way ANOVA revealed no significant effect of Clinical Group on the eye-motor switching rate, (*F*_3, 139_ = 1.67, *p* = .17, *η*² = 0.035). This result remained non-significant after controlling for differences between groups in age and gender via an ANCOVA (*F*_3, 137_ = 1.33, *p* = .27, *η*² = 0.03). As illustrated in Figure S1, eye-motor switching rates were numerically similar across groups. In contrast to H1a, the CHR-P and HC groups displayed the highest numerical means, while the FEP group showed the lowest switching rate. Our pre-registered comparison of HC versus CHR-P did not reveal a significant difference (*F*_1, 71_ < 1, *p* = .34, *η*² = 0.01), and it was also not significant when controlling for gender and age (*F*_1, 69_ < 1, *p* = .43, *η*² = 0.01). Furthermore, the dimensional analysis did not yield a significant correlation between the eye-motor switching rate and psychosis-risk symptom severity. (*r* = .028, *p* = .74). Thus, our pre-registered hypotheses H1a was not supported.

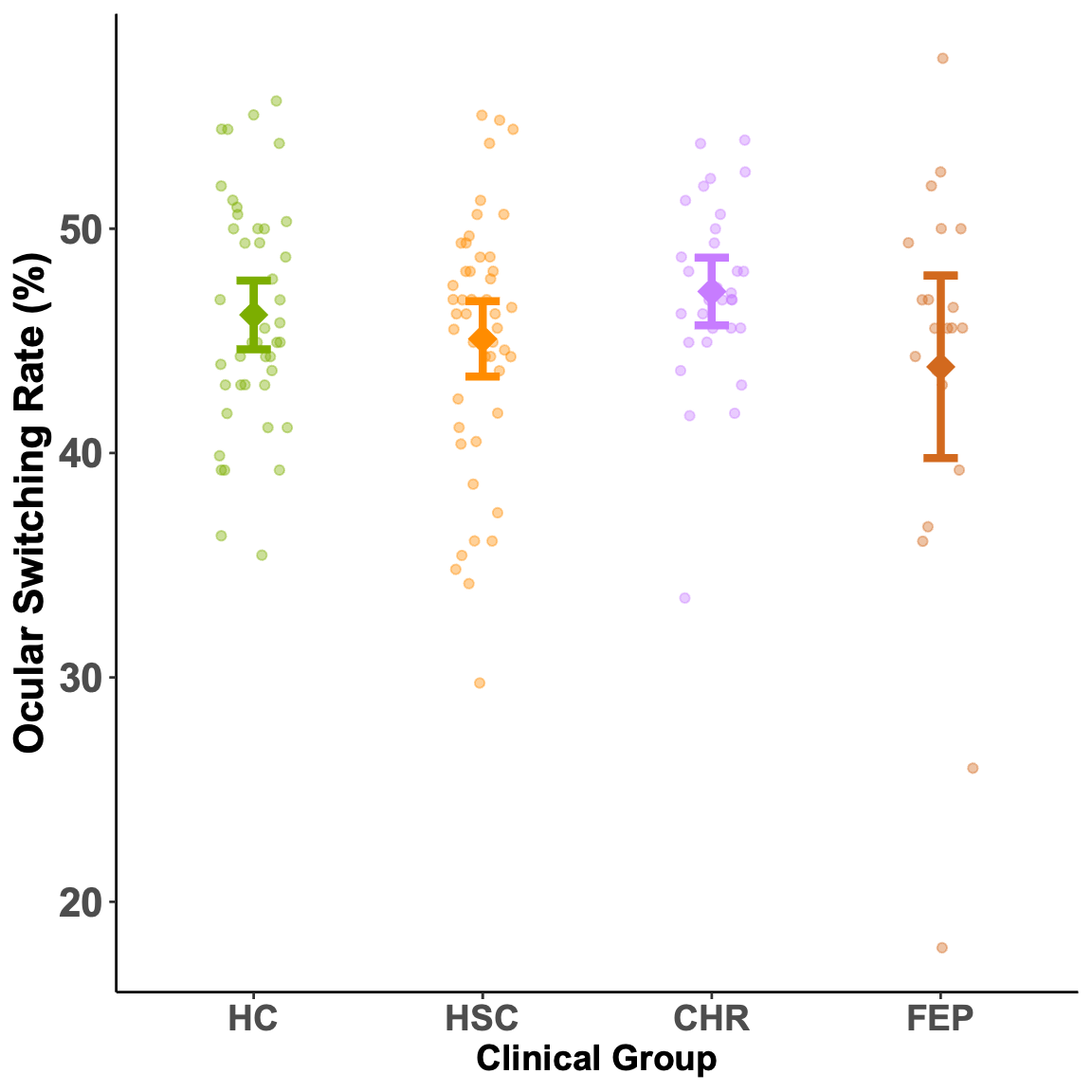

**Figure S3. Ocular motor switching rates across clinical groups.** Ocular motor switching rates, representing the frequency with which participants alternate their overall binary gaze direction on succeeding trials (i.e., right vs. left) regardless of the cue presented, were numerically similar across the psychosis continuum, with no significant differences observed between groups. Colored dots represent individual participants, while diamonds indicate group means and error bars represent 95% confidence intervals.

### 3.2.1 H1b: Group differences in model-based ocular motor perseverance

H1b predicted an increase in the model-based parameter (⍴) capturing ocular perseverance in the CHR-P group relative to HC. In addition, it predicted a significant positive correlation between ocular and increased psychosis-risk symptom severity. Contrary to our hypothesis, a one-way ANOVA revealed no significant effect of Clinical Group on the eye-motor switching rate, (*F*_3, 137_ = 1.22, *p* = .31, *η*² = 0.026). This result remained non-significant after controlling for differences between groups in age and gender via an ANCOVA (*F*_3, 135_ < 1, *p* = .44, *η*² = 0.02). Our pre-registered comparison of HC versus CHR-P did not reveal a significant difference (*F*_1, 73_ = 2.06 *p* = .16 *η*² = 0.03), and it was also not significant when controlling for gender and age (*F*_1, 71_ = 1, *p* = .20, *η*² = 0.02). Furthermore, contrary to our hypothesis the dimensional analysis yield a negative correlation between psychosis-risk symptoms and ocular perseverance that was not significant (*r* = -.13, *p* = .13). Thus, our pre-registered hypotheses H1b that model-based ocular perseverance is positively associated with psychosis-risk was not supported.

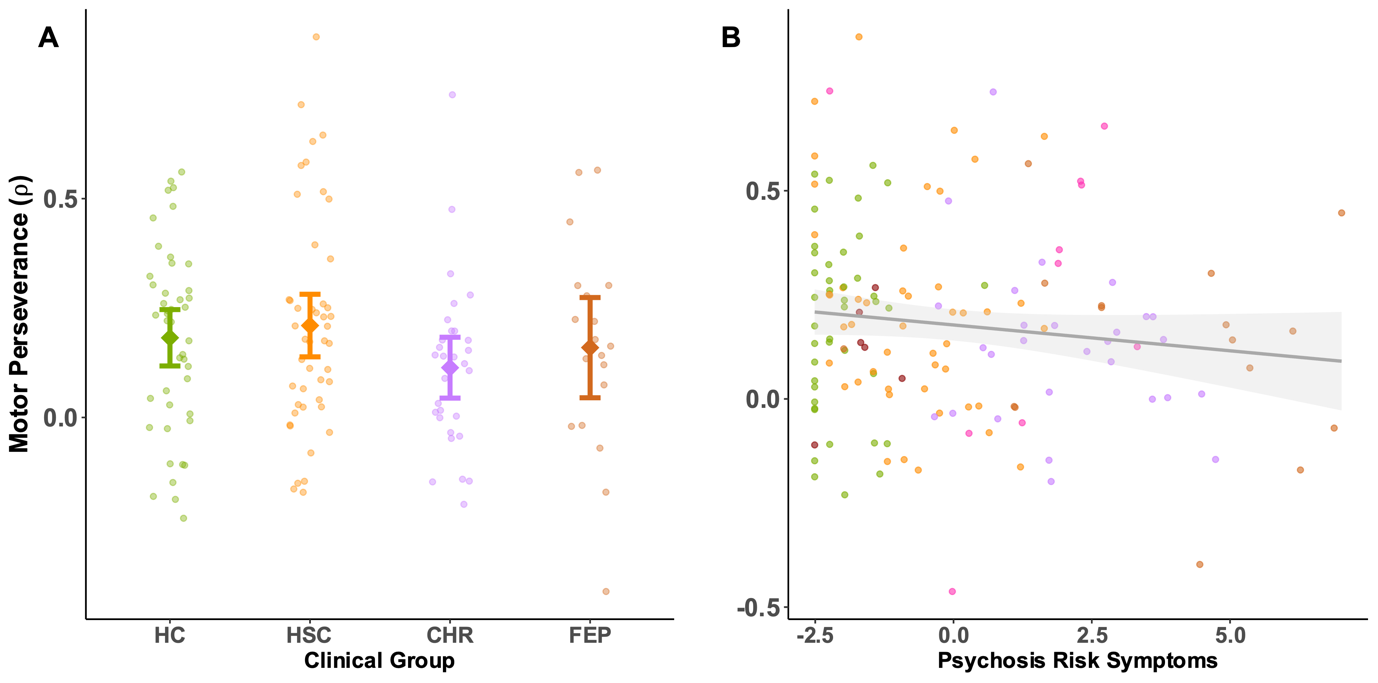

**Figure S4. Model-based ocular motor perseverance (⍴) by clinical group and symptom severity.** **(A) Group comparisons of motor perseverance.** No significant differences in motor perseverance were observed between clinical groups. The parameter ⍴ captures the tendency to repeat the same gaze direction on successive trials regardless of preceding feedback. Colored dots represent individual participants, while diamonds indicate group means and error bars represent 95% confidence intervals. **(B) Dimensional relationship with symptom severity.** There was no significant relationship between levels of motor perseverance and the severity of psychosis-risk symptoms. The solid line represents the linear regression fit with the shaded area indicating the 95% confidence interval; individual data points are colored by clinical group.

### 3.2.2 H2: Group differences in ocular learning mechanisms

Our second hypothesis (H2) proposed that ocular learning mechanisms are altered across the psychosis continuum. This was evaluated through three distinct sub-hypotheses focusing on trial-by-trial behavioral following the previous trial’s outcome and model-based parameters.

**3.2.2.1 H2a: Ocular learning following feedback across clinical groups**

First, H2a examined the sensitivity to previous trial’s outcome, predicting that the CHR-P group would demonstrate a reduced difference in ocular rule-switching following correct versus incorrect trials compared to HC. This was tested using a mixed logistic regression (*ocular rule switch ~ preceding trial accuracy * clinical group*). As expected, there was a significant main effect of preceding accuracy (*β* = -0.34, *z* = -6.87, *p* < .001), indicating that across both groups, participants were significantly more likely to switch their ocular rule following an incorrect trial than a correct trial. However, contrary to our pre-registration, the interaction between preceding accuracy and clinical group did not reach statistical significance (*β* = 0.14, *z* = 1.76, *p* = .079). However, the direction of the effect was numerically consistent with H2a, and t the difference in switching probability between correct and incorrect trials was smaller in the CHR-P group compared to the HC group. Finally. there was no significant main effect of clinical group on the overall switching rate (*β* = -0.04, *z* = -0.51, *p* = .61).

To examine the relationship between ocular learning and psychosis-proneness dimensionally across the full sample, we conducted a mixed-effects logistic regression using the standardized psychosis-spectrum symptoms’ Principal Component (SIPS PC) as a continuous measure of symptom severity. As expected, we observed a significant main effect of preceding accuracy (*β* = -0.27, *z* = -9.42, *p* < .001), confirming that participants generally switched more frequently following incorrect trials. Crucially, the interaction between preceding accuracy and the SIPS PC was significant (*β* = -0.08, *z* = -2.70, *p* = .007). However, contrary to our pre-registered hypothesis, the negative direction of this interaction coefficient indicates that elevated symptoms was associated with an *increased* difference in switching behavior between correct and incorrect trials. No significant main effect of the SIPS PC on overall switching rates was observed (*β* = 0.03, *z* = 0.95, *p* = .34).

In summary, our results provide divergent evidence regarding feedback sensitivity: while the categorical comparison between HC and CHR-P participants trended toward reduced sensitivity (H2a), the dimensional analysis across the full sample revealed an exaggerated behavioral adjustment. These divergent patterns preclude a definitive conclusion regarding the relationship between psychosis-proneness and trial-by-trial ocular learning within this specific cohort.

**3.2.2.2 H2b: Ocular accuracy across clinical groups**

H2b predicted reduced ocular rule accuracy in the CHR-P group relative to HC, alongside a significant negative relationship between accuracy and psychosis-proneness symptoms. Contrary to our hypothesis, a one-way ANOVA revealed no significant effect of Clinical Group on ocular rule accuracy (*F*_3, 139_ = 1.74, *p* = .16, *η*²= 0.036). This result remained non-significant after controlling for age and gender via an ANCOVA *F*_3, 137_ = 1.25, *p* = .29, *η*² = 0.027). As illustrated in Table S7, ocular rule accuracy followed a descending numerical trend across the psychosis continuum, being highest in the HC group and lowest in the FEP group. Despite this pattern, our pre-registered comparison of HC versus CHR-P did not reach statistical significance (*F*_1, 73_ = 1.39, *p* = .24, *η*²= 0.019). However, the dimensional analysis across the full sample revealed a marginal, however non-significant, negative correlation between ocular rule accuracy and psychosis-risk symptom severity (*r* = -0.17, *p* = .052). While this trend-level result is numerically consistent with the predicted direction of H2b, the categorical predictions for the CHR-P group were not supported.

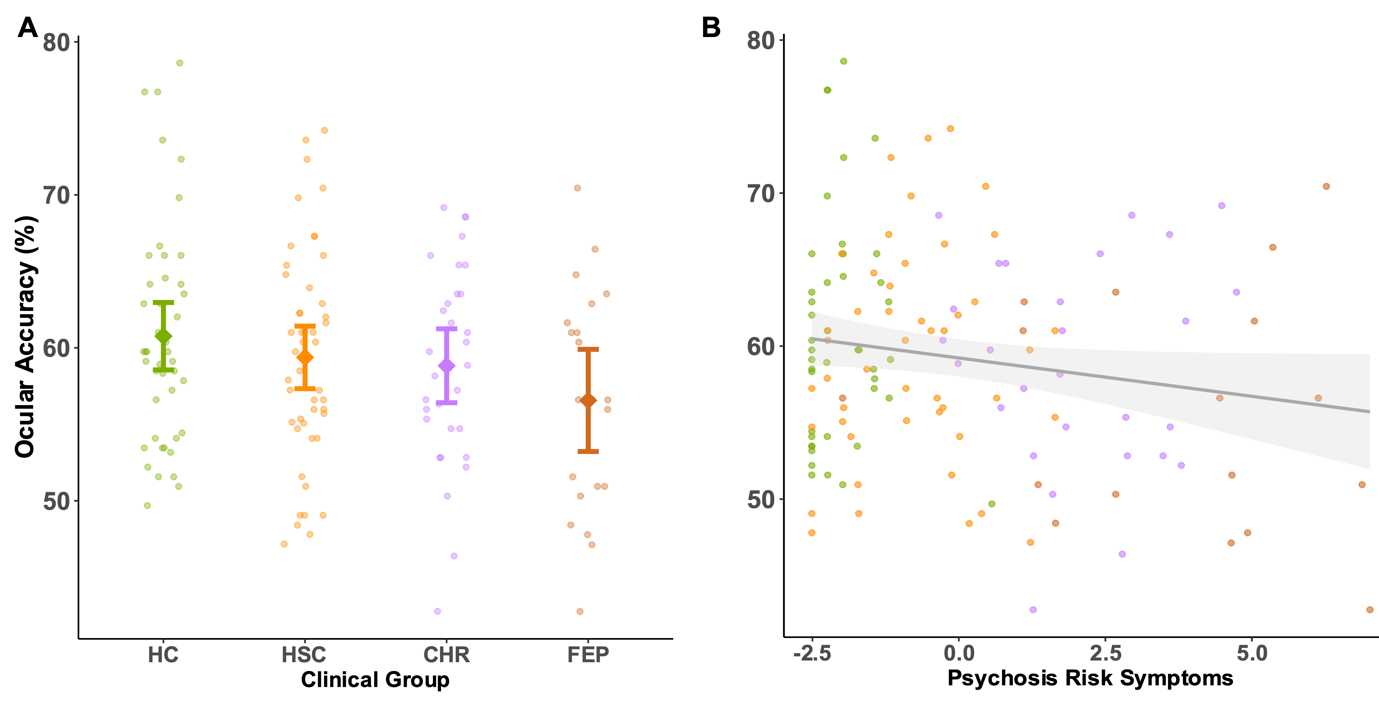

**Figure S5. Ocular rule accuracy by clinical group and symptom severity.** **(A) Group comparisons of ocular rule accuracy.** No significant differences in accuracy were observed between clinical groups, though a descending numerical trend was evident across the psychosis continuum, with the highest accuracy in the HC group and the lowest in the FEP group. Colored dots represent individual participants, while diamonds indicate group means and error bars indicate 95% confidence intervals. **(B) Dimensional relationship with symptom severity.** Analysis revealed a marginal, non-significant negative relationship between ocular rule accuracy and the severity of psychosis-risk symptoms. The solid line represents the linear regression fit with the shaded area indicating the 95% confidence interval; individual data points are colored by clinical group.

**3.2.2.3 H2c: Model-based stochasticity across clinical groups**

H2c predicted increased stochasticity (the epsilon parameter in the Win-Stay Lose-Switch model) in the CHR-P group relative to HC, alongside a significant positive relationship between epsilon values and psychosis-proneness symptoms. Contrary to our hypothesis, a one-way ANOVA revealed no significant effect of Clinical Group on the epsilon parameter (*F*_3, 139_ = 0.86, *p* = .47, *η*² = 0.018). This result remained non-significant after controlling for age and gender via an ANCOVA (*F*_3, 137_ = 0.46, *p* = .71, *η*² = 0.010). As illustrated in Table S7, epsilon values followed a non-significant increasing numerical trend across the psychosis continuum, with the FEP group exhibiting the highest numerical level of stochasticity. However, our pre-registered comparison of HC versus CHR-P did not reveal a significant difference (*F*_1, 73_ = 0.26, *p* = .61, *η*² = 0.004). Furthermore, the dimensional analysis across the full sample did not yield a significant correlation between the epsilon parameter and psychosis-risk symptom severity (*r* = 0.10, *p* = .26). Thus, our pre-registered hypothesis H2c was not supported.

In summary, our second hypothesis regarding altered ocular learning mechanisms across the psychosis continuum was largely not supported. While numerical trends in rule accuracy (H2b) and stochasticity (H2c) were generally consistent with a psychosis-continuum model, categorical comparisons between HC and CHR-P participants failed to reach statistical significance. Moreover, the trial-by-trial switching analysis (H2a) yielded divergent results between group-based and dimensional approaches. Collectively, these findings suggest that ocular learning mechanisms do not robustly differ between clinical groups.

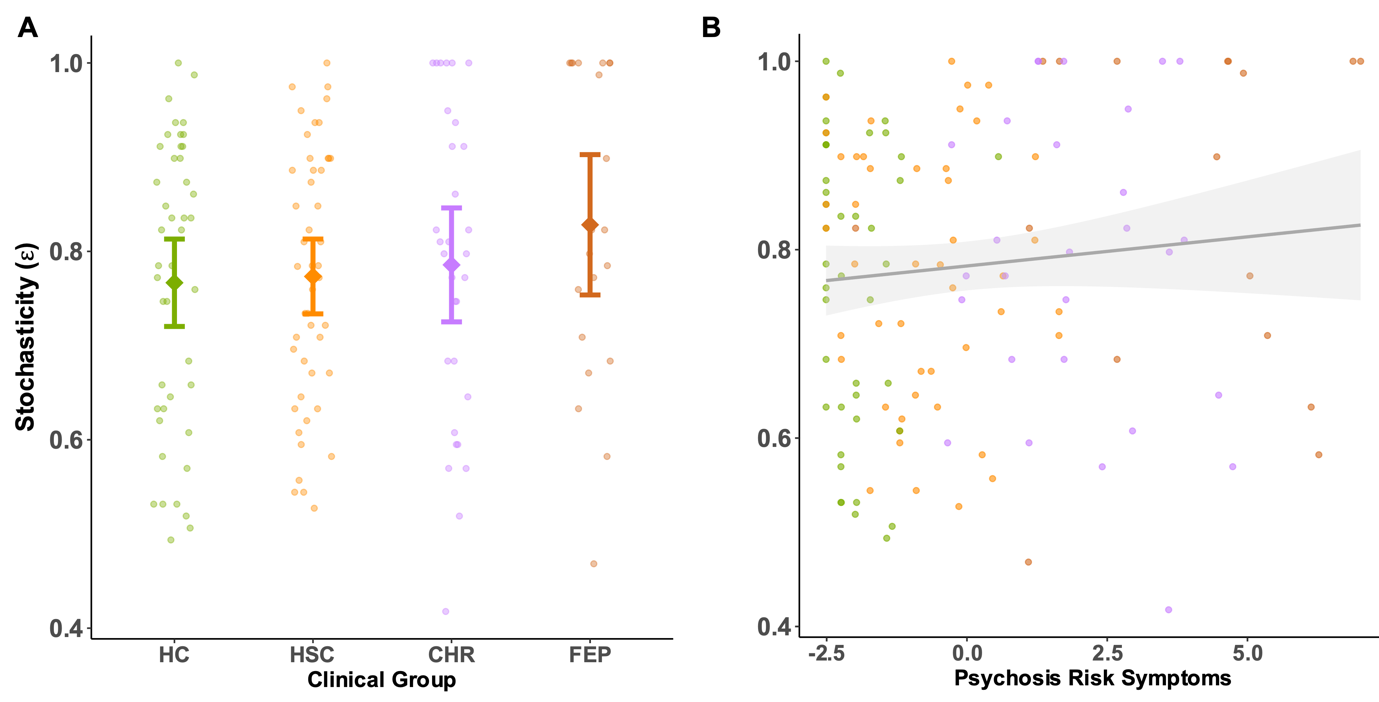

**Figure S6. Model-based stochasticity by clinical group and symptom severity.** **(A) Group comparisons of stochasticity.** No significant differences in stochasticity (i.e., ε) were observed between clinical groups, though values followed a non-significant increasing numerical trend across the psychosis continuum. The parameter ε represents the level of randomness or "noise" in trial-by-trial switching behavior. Colored dots represent individual participants, while diamonds indicate group means and error bars represent 95% confidence intervals. **(B) Dimensional relationship with symptom severity.** There was no significant relationship between levels of stochasticity and the severity of psychosis-risk symptoms. The solid line represents the linear regression fit with the shaded area indicating the 95% confidence interval; individual data points are colored by clinical group.

**3.2.3 H3: Global Metacognition across clinical groups**

H3 predicted a significant decrease in signed global metacognition for the CHR-P group relative to HC, reflecting an underestimation of explicit task performance. We further expected a significant negative correlation between global metacognition and psychosis-proneness symptoms. Contrary to our hypothesis, a one-way ANOVA revealed no significant effect of Clinical Group on global metacognition (*_F_*_3, 124_= 1.06, *p* = .37, *η*² = 0.025). This result remained non-significant after controlling for age and gender via an ANCOVA (*F*_3, 122_ = 1.26, *p* = .29, *η*² = 0.030).

As illustrated in Table SXX, the CHR-P group was the only group to numerically underestimate their performance, whereas the HC group showed a slight overestimation. However, our pre-registered comparison of HC versus CHR-P did not reach statistical significance (*F*_1, 67_ = 1.97, *p* = .17, *η*² = 0.029). Furthermore, the dimensional analysis revealed no significant correlation between global metacognition and psychosis-risk symptom severity (*r* = -0.04, *p* = .61). Thus, despite a numerical trend consistent with previous exploratory findings, our pre-registered hypothesis H3 was not supported in the current sample.

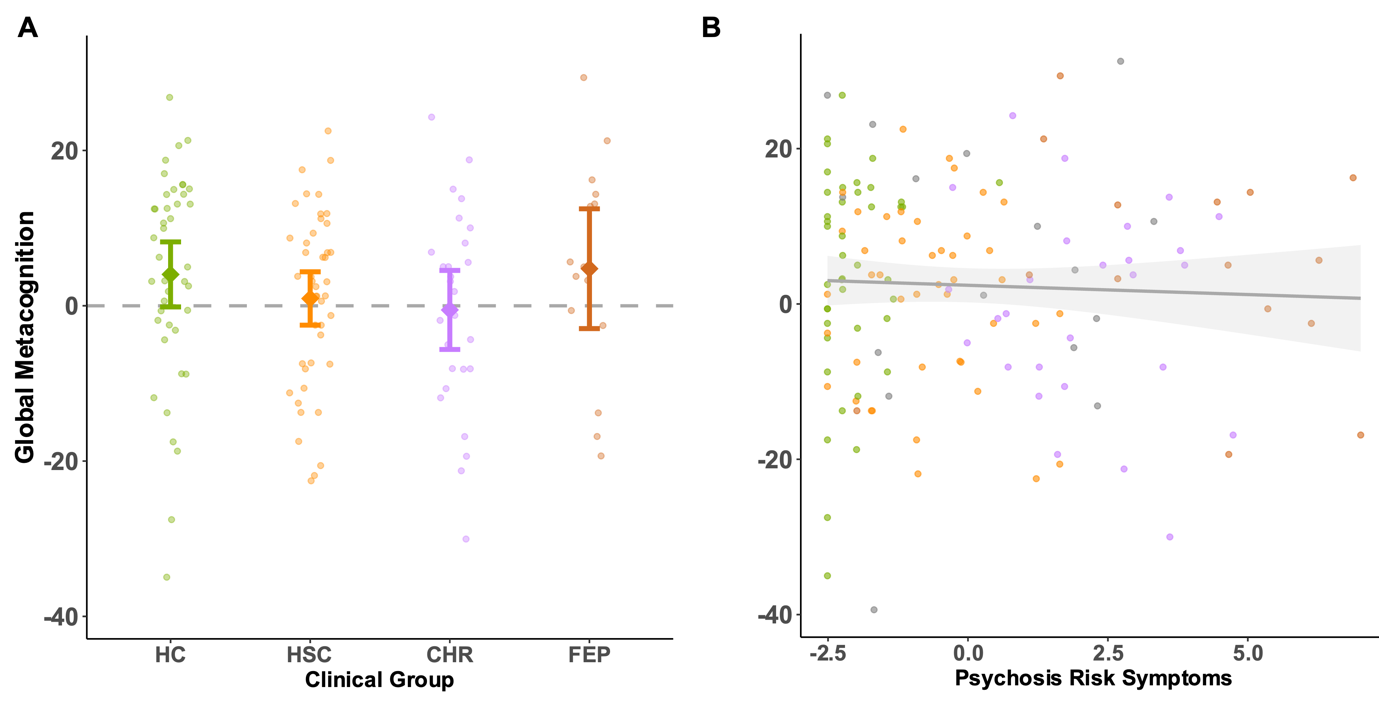

**Figure S7. Global metacognition by clinical group and symptom severity.** **(A) Group comparisons of global metacognition.** No significant differences in global metacognition were observed between clinical groups. Global metacognition reflects the signed difference between estimated and actual task performance, with negative values indicating an underestimation of performance and positive values an overestimation. The horizontal dashed line at zero represents accurate performance estimation. Colored dots represent individual participants, while diamonds indicate group means and error bars represent 95% confidence intervals.**(B) Dimensional relationship with symptom severity.** There was no significant relationship between global metacognition and the severity of psychosis-risk symptoms. The solid line represents the linear regression fit with the shaded area indicating the 95% confidence interval; individual data points are colored by clinical group.

| Hypothesis | Construct | HC | HSC | CHR | FEP |
| --- | --- | --- | --- | --- | --- |
| H1a | Ocular Motor Switching Rate  (%) | 46.15 [44.62, 47.69] | 45.09 [43.41, 46.76] | 47.20 [45.69, 48.71] | 43.85 [39.78, 47.91] |
| H1b | Motor Perseverance  (⍴) | 0.18 [0.12, 0.25] | 0.21 [0.14, 0.28] | 0.11 [0.04, 0.18] | 0.16 [0.05, 0.27] |
| H2b | Ocular Rule Accuracy  (%) | 60.76 [58.56, 62.96] | 59.37 [57.33, 61.42] | 58.83 [56.41, 61.24] | 56.56 [53.22, 59.90] |
| H2c | Model-based Stochasticity (ε) | 0.77 [0.72, 0.81] | 0.77 [0.73, 0.81] | 0.79 [0.73, 0.85] | 0.83 [0.75, 0.90] |
| H3 | Global Metacognition | 4.04 [-0.15, 8.23] | 0.94 [-2.50, 4.38] | -0.54 [-5.63, 4.55] | 4.78 [-2.94, 12.49] |

**Table S7.** Summary of Pre-registered Outcome Measures by Clinical Group (Mean [95% CI])
